# Characterization of Hydrophobic Interactions of SARS-CoV-2 and MERS-CoV Spike Protein Fusion Peptides Using Single Molecule Force Measurements

**DOI:** 10.1101/2022.03.05.483104

**Authors:** Cindy Qiu, Gary R. Whittaker, Samuel H. Gellman, Susan Daniel, Nicholas L. Abbott

## Abstract

We address the challenge of understanding how hydrophobic interactions are encoded by fusion peptide sequences within coronavirus (CoV) spike proteins. Within the fusion peptides of SARS-CoV-2 and MERS-CoV, a largely conserved peptide sequence called FP1 (SFIEDLLFNK and SAIEDLLFDK in SARS-2 and MERS, respectively) has been proposed to play a key role in encoding hydrophobic interactions that drive viral-host cell membrane fusion. While a non-polar triad (LLF) is common to both FP1 sequences, and thought to dominate the encoding of hydrophobic interactions, FP1 from SARS and MERS differ in two residues (Phe 2 versus Ala 2 and Asn 9 versus Asp 9, respectively). Here we explore if single molecule force measurements can quantify hydrophobic interactions encoded by FP1 sequences, and then ask if sequence variations between FP1 from SARS and MERS lead to significant differences in hydrophobic interactions. We find that both SARS-2 and MERS wild-type FP1 generate measurable hydrophobic interactions at the single molecule level, but that SARS-2 FP1 encodes a substantially stronger hydrophobic interaction than its MERS counterpart (1.91 ± 0.03 nN versus 0.68 ± 0.03 nN, respectively). By performing force measurements with FP1 sequences with single amino acid substitutions, we determine that a single residue mutation (Phe 2 versus Ala 2) causes the almost threefold difference in the hydrophobic interaction strength generated by the FP1 of SARS-2 versus MERS, despite the presence of LLF in both sequences. Infrared spectroscopy and circular dichroism measurements support the proposal that the outsized influence of Phe 2 versus Ala 2 on the hydrophobic interaction arises from variation in the secondary structure adopted by FP1. Overall, these insights reveal how single residue diversity in viral fusion peptides, including FP1 of SARS-CoV-2 and MERS-CoV, can lead to substantial changes in intermolecular interactions proposed to play a key role in viral fusion, and hint at strategies for regulating hydrophobic interactions of peptides in a range of contexts.

**SIGNIFICANCE:** Fusion of coronaviruses (CoVs) and host cells is mediated by the insertion of the fusion peptide (FP) of the viral spike protein into the host cell membrane. Hydrophobic interactions between FPs with their host cell membranes regulate the viral membrane fusion process and are key to determining infection ability. However, it is not fully understood how the amino acid sequences in FPs mediate hydrophobic interactions. We use single-molecule force measurements to characterize hydrophobic interactions of FPs from SARS-CoV-2 and MERS-CoV. Our findings provide insight into the mechanisms by which the amino acid composition of FPs encodes hydrophobic interactions and their implications for fusion activity critical to the spread of infection.

## INTRODUCTION

The ongoing coronavirus (CoV) outbreak involving SARS-CoV-2 has galvanized efforts to understand the biophysical interactions between coronaviruses and host cells as a foundation for developing novel therapeutics, new diagnostic tools, and the ability to predict future outbreaks (1-5). Prior outbreaks of coronaviruses include severe acute respiratory syndrome (SARS-CoV) and Middle East respiratory syndrome (MERS-CoV) (6,7).

The spike (S) glycoprotein present in the envelope of SARS-CoV-2 plays a key role in regulating viral entry into host cells (8). The S protein encompasses subunits S1, a receptor binding domain (RBD), and S2, which is directly involved in the cell membrane fusion event (Figure 1a). The receptor binding domain identifies and binds to a receptor on the host cell, e.g. ACE2 for SARS-CoV-2, which is followed by fusion of the viral and host cell membranes to enable viral entry into the host cell. Coronaviruses have flexible entry pathways and can fuse with host cells at the plasma membrane or in the endosome; evidence for either routes exists for SARS-CoV-2, which may be based on the variant and cell type infected (9,10). The process of selecting between the two routes is believed to be triggered by the availability of proteases in the surrounding environment, as shown with MERS-CoV (11). After cleavage by proteases, the S protein undergoes conformational changes to expose the fusion peptide (FP) of S2 for interaction with the host cell (8). Cell membrane fusion is further driven by association of two heptad repeat domains within S2 into a six-bundle assembly, one closer to the N-terminus (HRN) and another towards the C-terminus (HRC) (Figure 1a) (12,13). Important to our study, a crucial initial step for successful membrane fusion and subsequent infection is insertion of the FP within S2 into the host cell membrane to trigger the fusion event (14).

**Figure 1.**
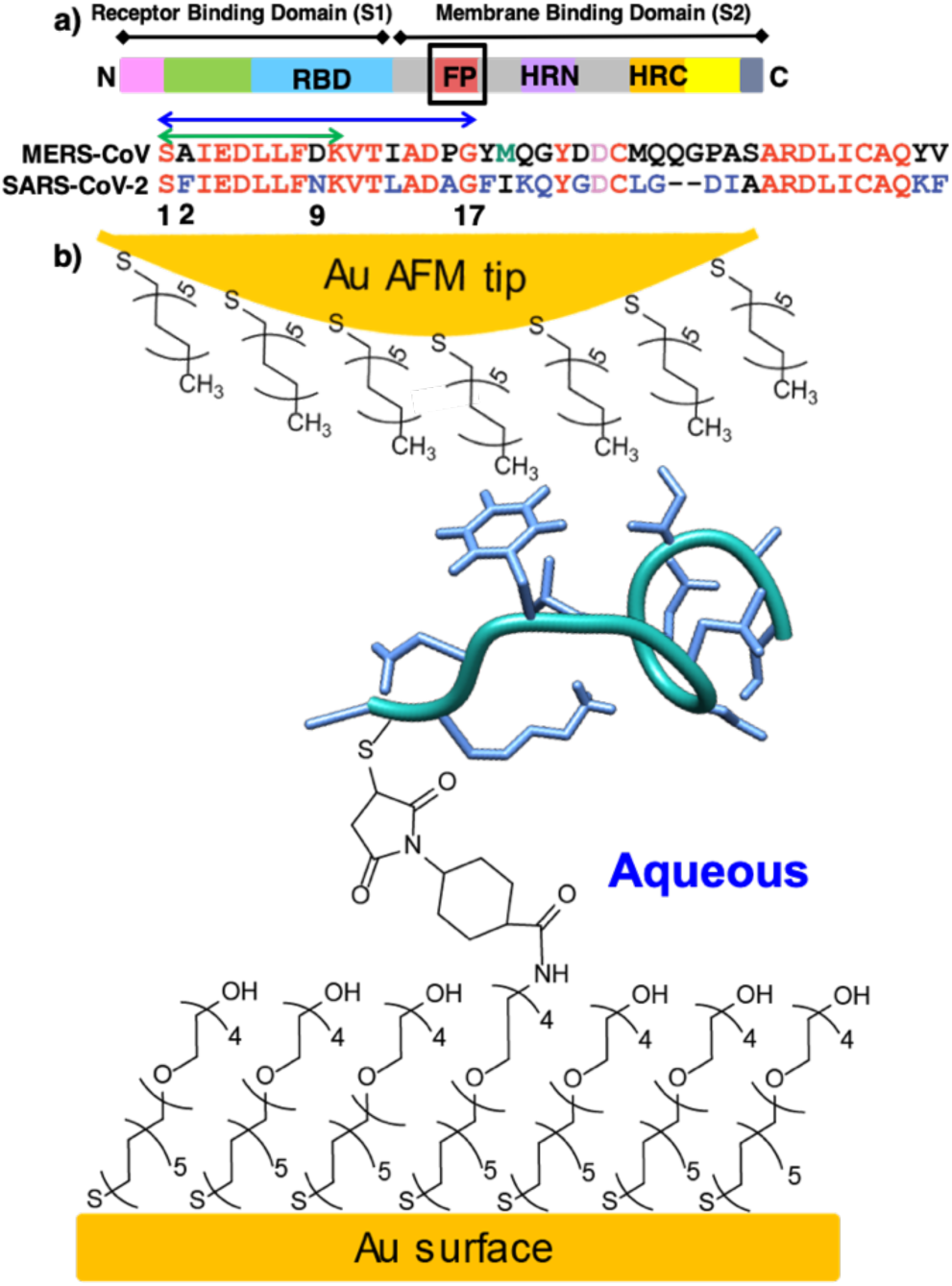
a) Schematic representation of functional elements of the coronavirus (CoV) spike (S) protein. The green arrow denotes the FP1 portion of the FP used in the first part of our study described in the text (labeled S1 to K10 in SARS-CoV-2 and MERS-CoV). The blue arrow indicates the SARS-CoV-2 FP sequence used in the second part of our study (S1 to G17). b) Schematic representation of an alkyl-terminated AFM tip interacting with single FP1 molecule covalently immobilized to a chemically modified gold surface in aqueous solution.

Recent efforts have succeeded in advancing our understanding of the structural components of the S protein using techniques such as cryo-electron microscopy and X-ray crystallography (15-18). These studies include characterization of the interactions of S1 of the S protein with ACE2 (19-22). In contrast, the current understanding of the interactions of the FP is limited. Within the coronavirus family, the FP was first identified for SARS-CoV based on its high degree of sequence homology across *Coronaviridae*, and shown to be present as two domains encompassing 15-42 largely conserved non-polar and charged amino acids (23,24). Subsequent studies have suggested that within the FP, two functionally distinct regions adjacent to each other, “FP1” and “FP2,” cooperate to form a bipartite fusion platform (25) that encodes hydrophobic (lipid-binding) and ionic (i.e. Ca^2+^ binding) interactions to promote fusion (25-29). In particular, mutagenesis studies have identified a non-polar motif within the FP1 comprising Leu-Leu-Phe (LLF) to play a key role in membrane fusion (25,27,29,30). The non-polar nature of the motif is thought to promote hydrophobic interactions with cell membranes (31-33), although direct experimental characterization of the interactions encoded by the sequence of amino acids in FP1 has not been explored.

The hydrophobic interaction is a water-mediated attraction between non-polar domains of relevance to an array of biological contexts, from protein folding to assembly of membranes (34-37). In these various contexts, the non-polar domains do not occur in isolation, but rather, are found proximal to polar and charged groups, forming nanoscopic chemical patterns (38). Simulations have suggested that hydrophobic interactions cannot simply be described as an additive consequence of the functional groups present, but rather, depend on the context in which these functional group are placed (38-42). However, experimental studies of this phenomenon remain challenging.

Recent studies by our group and others have established a methodology that uses an Atomic Force Microscope (AFM) to measure hydrophobic interactions between model β-oligopeptides and non-polar surfaces at the single molecule level (43-47). These studies have uncovered context-dependent hydrophobic interactions mediated by the three-dimensional nano-scale patterning of polar and charged functional groups placed proximal to non-polar moieties (45). Specifically, hydrophobic interactions encoded by conserved non-polar domains were found to be strongly modulated by the identity of adjacent charged and polar moieties (43-45). In our current study, we leverage this approach and understanding of the origins of hydrophobic interactions to characterize the hydrophobic interactions of FP sequences of SARS-CoV-2 and MERS-CoV (Figure 1b). We identify the hydrophobic interactions encoded by the FP sequences by allowing them to interact with model non-polar surfaces. The non-polar surfaces are not intended as models of biological interfaces but rather serve as a reference surface with which to unmask the effects of amino acid identity on hydrophobic interactions encoded by FP sequences (48-50). This foundational knowledge is necessary to guide the interpretation of future studies, including experiments that explore interactions between CoV FPs and mammalian cell membranes.

In the first part of this paper, we examine the interactions of a undecamer (S1 to K10) from the FP1 domain of SARS-CoV-2 and MERS-CoV (Figure 1a, green arrow). These 11-amino acid sequences were selected for our initial studies as they include LLF but also encompass key differences between SARS-CoV-2 and MERS-CoV (23,25,27,29,51). Below we refer to these sequences as “SARS-2 FP1” and “MERS FP1.” SARS-2 FP1 contains Phe 2 and Asn 9, while these amino acids are replaced by Ala 2 and Asp 9, respectively, in MERS FP1. We set out to address two key questions: 1) Do SARS-2 FP1 and MERS FP1 sequences (S1 to K10, both including LLF) encode measurable hydrophobic interactions? 2) Do we measure a difference in hydrophobic interactions between the two sequences, thus revealing that the identity of the amino acids flanking LLF plays a key role in encoding hydrophobic interactions?

In the second part of our paper, we extend the length of the SARS-CoV-2 peptide sequence used in our study to encompass FP1 and a portion of the FP2 region (S1 to G17, Figure 1), enabling us to evaluate the generality of our findings from the first part of this study (Figure 1a, blue arrow). Below we refer to this extended sequence as “SARS-2 FP1+.” In particular, we probe the importance of amino acid identity in position 2 of this FP sequence by comparing the hydrophobic interactions generated by the wild-type sequence with a sequence of the same length in which Phe 2 was replaced by Ala 2 (Figure 4a, b). Additionally, given the role proposed for LLF in promoting membrane fusion in past studies (25,27,29,30), we examine the role of LLF in encoding hydrophobic interactions with a non-polar AFM tip. Specifically, we substitute LLF residues for Tyr, Ala, and Ser (see below for details). This set of experiments probes a key question: How important is LLF in encoding hydrophobic interactions of the fusion peptide of SARS-CoV-2? As detailed below, a key finding of our study is that single amino acid differences between FPs from SARS-CoV-2 and MERS-CoV can lead to large changes in the hydrophobic interactions encoded by the sequences. To provide insight into the mechanisms by which single amino acid substitutions can lead to such large changes in the strength of hydrophobic interactions, we characterize the conformations of FPs using circular dichroism (CD) and attenuated total reflectance – Fourier transform infrared spectroscopy (ATR-FTIR), the latter on surfaces similar to those used in our AFM studies.

## MATERIALS AND METHODS

### Materials

Tetraethylene glycol thiols terminated in hydroxyl (EG4) or amine groups (EG4N) were purchased from Prochimia (Gdynia, Poland). 1-Dodecanethiol (C12SH, 98%), triethanolamine HCl (TEA, 99%), phosphate-buffered saline (PBS), 1X with calcium and magnesium (Corning, NY; see Table S3 for composition of buffer), methanol (anhydrous, 99.8%), and ethanol (reagent, anhydrous, denatured) for preparing thiol solutions were purchased from Sigma-Aldrich (Milwaukee, WI). Sulfosuccinimidyl-4-(N-maleimidomethyl) cyclohexane-1-carboxylate (SSMCC) was purchased from ThermoFisher Scientific (Rockford, IL). All fusion peptides were purchased from Biomatik (Wilmington, DE). Ethanol (anhydrous, 200 proof) used for rinsing was purchased from Decon Labs (King of Prussia, PA). All chemicals were used without additional purification.

The AFM tips (triangular shaped, nominal spring constant of 0.01 N/m) were purchased from Bruker Nano (Camarillo, CA). Silicon wafers were purchased from Silicon Sense (Nashua, NH). A 45° multi-reflection germanium crystal and ATR-FTIR accessory were purchased from Pike Technologies (Madison, WI).

### Preparation of fusion peptide-decorated surfaces

Fusion peptides were synthesized via solid-phase methods by Biomatik and immobilized as detailed previously (43-47). In brief, we immobilized the fusion peptides onto mixed monolayers terminated in tetraethylene glycol (EG4) or aminotetraethylene glycol (EG4N) groups, using a mole fraction of EG4N of 0.002 to achieve a low surface density of immobilized peptides. This approach allowed us to measure adhesive interactions between single fusion peptide molecules immobilized onto mixed monolayers and the tip of the AFM (44-46,52). Previous work has shown that EG4N/EG4 mixed monolayers do not generate measurable adhesive forces with non-polar AFM tips in aqueous PBS buffer (52).

### Preparation of chemically functionalized AFM tips

Triangular-shaped cantilevers with nominal spring constants of 0.01 N/m were used for experiments involving fusion peptides. AFM tips were coated with a 2 nm layer of titanium and a 20 nm layer of gold by physical vapor deposition using an electron beam evaporator (44-46,52). Subsequently, the gold-coated tips were immersed in a 1 mM ethanolic solution of C12SH and incubated overnight for 18 hours. Upon removal from solution, tips were rinsed with ethanol, dried with a gentle stream of nitrogen, and directly transferred to the AFM fluid cell.

### AFM force measurements

Adhesion force measurements were performed using a Nanoscope IIIa Multitude AFM equipped with a fluid cell (Veeco Metrology Group, Santa Barbara, CA). Triangular-shaped silicon nitride cantilevers were used and functionalized as described above. The nominal spring constant of AFM tips used was 0.01 N/m. The spring constants of the cantilevers were calibrated using the thermal tuning method on the Asylum MFP-3D (Santa Barbara, CA) and determined to be 0.027 ± 0.003 N/m. Force measurements were performed at room temperature. Force curves were recorded using a constant contact time of 500 ms and retraction and approach speeds of 1,000 nm/s. Measurements of fusion peptides were performed in aqueous PBS. A “J” type scanner was used for the force measurements.

### ATR-FTIR measurements

Model non-polar monolayers and peptide monolayers were formed on gold-coated germanium ATR-FTIR crystal. First, the Ge crystal was coated with a 2 nm layer of gold by physical vapor deposition using an electron beam evaporator. Subsequently, the gold-coated crystal was immersed in a 1 mM ethanolic solution of C12SH and incubated overnight for 18 hours to create a non-polar monolayer mimicking the composition of our non-polar AFM tips. Upon removal from solution, the Ge crystal were rinsed with ethanol, and dried with a gentle stream of nitrogen. Next, this non-polar monolayer was incubated in 100 μM fusion peptide solution in DMSO. ATR-FTIR spectra were collected using a Horizontal Attenuated Total Reflectance (HATR) accessory from Pike Technologies paired with Nicolet iS50 FTIR Spectrometer from ThermoFisher Scientific. Each spectrum was acquired with a minimum of 300 scans with a resolution of 4 cm^-1^. Triplicate measurements were averaged for each peptide.

### Circular Dichroism (CD) measurements

CD spectroscopy measurements were performed using a JASCO J-1500 Spectrophotometer with 10 mm path length quartz cuvettes. All FPs were dissolved in PBS buffer containing 0.9 mM CaCl_2_ and 60 vol % MeOH in PBS to a 0.1 mM solution. CD spectra were collected at 25°C from 260 to 195 nm. Three independent sample solutions of each FP were prepared and measured. Blank spectra of PBS and 60 vol % MeOH were subtracted from the respective FP solution type. We estimated the percent helicity of 11-amino acid FP1 sequences in PBS (Figure 6a and Supporting Information Figure S7) to assess potential differences between sequences containing Phe 2 vs Ala 2 in bulk solution. First, we converted ellipticity (units of mdeg) to mean residue ellipticity (units of deg cm^2^ decimol^-1^) (53). Percent helicity was estimated as 100([*θ*]_222_ / -39,500 x (1 – 2.57/*n*)), where -39,500 represents the maximum theoretical mean residue ellipticity for a helix of *n* residues at 222 nm (54).

### Statistical Analysis

Adhesion force measurements performed to generate the histograms in Figures 2-4 are the averages of 6 independent samples (See Supporting Information Figure S1). Each histogram represents over 3,000 pull-off force curves (Supporting Information Figure S1). We performed *t*-tests to probe the statistical significance of differences in mean adhesion forces that we measured.

**Figure 2.**
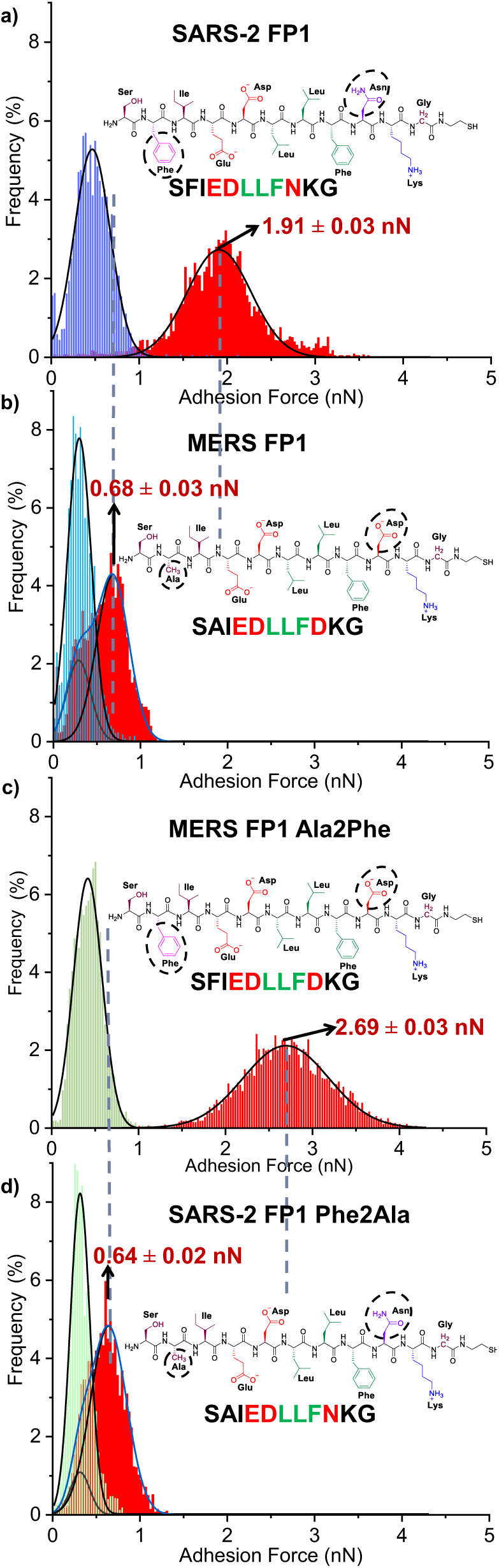
a) Adhesion forces of SARS-2 FP1 in aqueous buffer (PBS) containing 0.9mM Ca^2+^ attributed to hydrophobic interactions (red histogram). The purple histogram shows adhesion forces after addition of 60 % MeOH. b) Adhesion forces of MERS FP1 in aqueous buffer (PBS) containing 0.9mM Ca^2+^ attributed to hydrophobic interactions (red histogram). The blue histogram shows adhesion forces after addition of 60 % MeOH. c) Adhesion forces of MERS FP1 Ala2Phe in aqueous buffer (PBS) containing 0.9mM Ca^2+^ attributed to hydrophobic interactions (red histogram). The green histogram shows adhesion forces after addition of 60 % MeOH. h) Adhesion forces of Phe with Ala in aqueous buffer (PBS) containing 0.9mM Ca^2+^ attributed to hydrophobic interactions (red histogram). The green histogram shows adhesion forces after addition of 60 % MeOH. Adhesion force histograms were obtained using over 3,000 pull-off force curves from 6 independent samples. Data show mean ± s.e.m.

## RESULTS AND DISCUSSION

We began by measuring the adhesive pull-off forces between SARS-2 FP1 or MERS FP1 sequences and a model non-polar surface, with the goal of characterizing the hydrophobic interactions encoded by the peptides. Following a methodology described previously, we measured the adhesion force between single surface-immobilized FPs (immobilized via a terminal SH group) and a non-polar AFM tip in either aqueous PBS buffer, or PBS to which 60 vol % methanol was added (43-47). The PBS used in our measurements contained Ca^2+^, as prior experiments have demonstrated the ability of the ion to promote membrane fusion with MERS-CoV and SARS-CoV (25,28). Previously, we established that addition of 60 vol % methanol to aqueous buffer eliminates a majority of hydrophobic interactions mediated by non-polar domains without measurably disrupting ionic (i.e., electrical double layer interactions) and van der Waals interactions (44-50,55). Correspondingly, adhesion forces measured in 60 vol % MeOH/40 vol % PBS buffer are largely generated by van der Waals and electrical double layer interactions. This methodology was applied previously to conformationally stable β-peptide oligomers that form helices, which provides predictable presentation of non-polar and polar residues under various solution conditions. In contrast, the FPs used in our current study are oligomers of α-amino acids that lack the conformational rigidity of β-peptides. Accordingly, the secondary structure and thus spatial presentation of the residues of FP1 depend on solution environment and interaction with interfaces. Below we return to this important consideration in the context of CD and ATR-FTIR measurements that characterize the conformations of FP1 in bulk solution and in contact with non-polar surfaces.

We measured the mean pull-off force of SARS-2 FP1 in PBS to be 1.91 ± 0.03 nN (Figure 2a, red histogram). Upon addition of 60 vol % MeOH to PBS, the adhesion force decreased to 0.47 ± 0.01 nN (Figure 2a, blue histogram). As there was no overlap in the histograms of pull-off forces measured in PBS or PBS containing 60 vol % MeOH, we interpret the pull-off forces measured in PBS buffer (red data) to be dominated by hydrophobic interactions, while the forces in 60 vol % MeOH (purple data) correspond largely to van der Waals and electrical double layer interactions (44,45). In *t*-tests that we performed to assess statistical significance, with the exception of one comparison described below, we found p-values for all comparisons we make to be < 0.05, revealing the presence of a statistical difference at a significance level of 95%. (See Supporting Information Tables S1 and S2).

To establish that the pull-off forces reported in Figure 2a are the result of interactions between the non-polar tip of the AFM and a single peptide molecule, we performed two sets of control experiments. First, we measured pull-off forces using a SSMCC-activated monolayer that was treated with β-mercaptoethanol before incubation with thiol-terminated FP sequences (see Supporting Information Figure S1). The thiol group of β-mercaptoethanol reacts with the maleimide group of SSMCC, thereby preventing covalent attachment of FPs. We found that measurements performed with the β-mercaptoethanol-treated surfaces and the AFM tip led to largely non-adhesive events (See Supporting Information, Figure S1) (43).

Second, we measured adhesion forces between the non-polar AFM tip and FP-decorated monolayers in which aminotetraethylene glycol (EG4N) groups were reduced to a mole fraction 0.001, thus lowering the number density of surface-immobilized peptides (see Materials and Methods). This procedure was predicted to lower the frequency of adhesive events but not change the magnitude of adhesion forces, relative to measurements obtained with a 0.002 mole fraction of EG4N. Our measurements were consistent with this prediction (see Figure 2; Supporting Information, Figure S2). Overall, these two control experiments provide support for our conclusion that the adhesion forces reported in Figure 2 result from interactions of the non-polar AFM tip and FPs at the single molecule level.

Next, we next performed measurements with the MERS FP1 sequence in PBS and 60 vol % MeOH added to PBS (Figure 2b). As detailed elsewhere (44,45), because the histograms of pull-off forces measured in the two solvent environments partially overlap, we identified the hydrophobic contribution to the pull-off force measured in PBS by fitting two Gaussian distributions, one of which was based on the same distribution as that measured in 60 vol % MeOH (Figure 2b) (44,45) (see above for additional discussion of this methodology). From this analysis, we conclude that the MERS FP1 sequence generated a mean hydrophobic force in PBS of 0.68 ± 0.03 nN (Figure 2b, red histogram). When 60 vol % MeOH was added to buffer, the mean pull-off force decreased to 0.29 ± 0.01 nN (Figure 2b, blue histogram).

Here, we make two preliminary observations by comparing the pull-off forces measured using the SARS-2 FP1 and MERS FP1 sequences (Figure 2a, b). First, we observe that the forces measured in 60 vol % MeOH/40 vol % PBS are comparable for the two sequences. This suggests that the two FP1 sequences encode similar van der Waals and electrical double layer interactions with the non-polar tip of the AFM. Second, and more significantly, our measurements suggest that the SARS-2 FP1 sequence encodes a substantially stronger hydrophobic interaction than its MERS FP1 counterpart (1.91 ± 0.03 versus 0.68 ± 0.03 nN, respectively). This result is an interesting one because both sequences possess the LLF non-polar triad, which previously has been proposed to dominate hydrophobic interactions of the FP1 (25,27,29,30). Our result hints that differences in the identity of amino acids flanking LLF in the FP1 from SARS-2 and MERS can regulate the strength of the hydrophobic interaction encoded by LLF by a factor of ∼3.

### Why are the hydrophobic interactions of FP1 from SARS-2 and MERS different?

The compositions of FP1 from SARS-2 FP1 and MERS FP1 differ at two positions in the sequence: Phe 2 vs. Ala 2 and Asn 9 vs. Asp 9, respectively (Figure 1a). To evaluate the role of these residues flanking LLF in encoding FP1 hydrophobic interactions, we performed force measurements with two sequences containing single point mutations (Figure 2c, d). The first mutated sequence replaced Ala 2 of MERS FP1 with Phe, while conserving Asp 9 (Figure 2c). Below we refer to this mutation as “MERS FP1 Ala2Phe.” The second mutated sequence replaced Phe 2 of SARS-2 FP1 with Ala while preserving Asn 9. Below this sequence is called “SARS-2 FP1 Phe2Ala” (Figure 2d).

We measured the mean pull-off force of MERS FP1 Ala2Phe in PBS to be 2.69 ± 0.03 nN (Figure 2c, red histogram). Upon addition of 60 vol % MeOH to PBS, the adhesion force decreased to 0.30 ± 0.01 nN (Figure 2c, green histogram). Similar to the SARS-2 FP1 wild-type sequence, the lack of overlap between the two histograms indicates that the pull-off forces in PBS are primarily hydrophobic in nature. In contrast, the mean hydrophobic pull-off force of SARS-2 FP1 Phe2Ala was only 0.64 ± 0.02 nN (Figure 2d, red histogram). When these measurements are combined with the results obtained using SARS-2 FP1 and MERS FP1 sequences (Figure 2a, b), we observe a correlation between the strength of hydrophobic interaction encoded by the sequence and the identities of the residues flanking LLF. In particular, the FP1 sequences with Phe 2 encode a hydrophobic interaction strength of 1.91 ± 0.03 or 2.69 ± 0.03 nN (with Asn 9 or Asp 9; Figure 2a, c, respectively). On the other hand, FP1 sequences with Ala leads to hydrophobic interactions of 0.68 ± 0.03 or 0.64 ± 0.02 nN (Asn 9 or Asp 9; Figure 2b and d, respectively). Significantly, FP1 sequences containing Phe 2 (Figure 2a and c) exhibit hydrophobic interactions that are substantially larger than sequences containing Ala 2 (Figure 2b and d).

The Phe 2-containing FP1 sequences comprise five non-polar amino acids (i.e., two Phe, one Ile, and two Leu). In contrast, the Ala 2-containing FP1 sequences contain four non-polar residues (one Phe, one Ile, two Leu). Interestingly, the results above reveal that the addition of a single non-polar amino acid to the FP1 sequence (i.e., increasing the number of non-polar amino acids in the sequence from 4 to 5; Ala 2 to Phe 2) can lead to a 3-4 fold increase in the strength of the hydrophobic interaction (from 0.64 ± 0.02 or 0.68 ± 0.03 to 1.91 ± 0.03 or 2.69 ± 0.03 nN). Overall, we conclude that the identity of the amino acid at position 2 (Ala 2 versus Phe 2) has an outsized influence on the magnitude of the hydrophobic interaction encoded by the FP1 sequences. In contrast, the mutations involving Asn 9 vs. Asp 9 have only a modest impact on the strength of the hydrophobic interaction.

Our insight above regarding the outsized role of Phe 2 in encoding the hydrophobic interactions of FP1 of SAR-CoV-2 are based on measurements performed with a peptide sequence comprising 11 amino acids. To explore the impact of Phe 2 on interactions encoded by longer sequences of amino acids from the FP of SARS-CoV-2, we next examined a 17-amino acid SARS-2 FP sequence that included a portion of FP2 (Figure 3). The six additional amino acid residues include two non-polar residues (Val and Leu). We refer to this sequence as “SARS-2 FP1+.” We compared the hydrophobic interactions encoded by SARS-2 FP1+ (Figure 3a) with a sequence with a single residue mutation, Phe2Ala (Figure 3b), which we call “SARS-2 FP1+ Phe2Ala”. Inspection of Figure 3a reveals that the 17-amino acid wild-type FP sequence in PBS encoded a hydrophobic pull-off force of 2.21 ± 0.02 nN (Figure 3a, red histogram). After 60 vol % MeOH was added to the PBS, the mean adhesion forces diminished to 0.33 ± 0.01 nN (Figure 3a, green histogram). When compared to SARS-2 FP1, which generated a hydrophobic interaction of 1.91 ± 0.03 nN (Figure 2a), we find that the additional six amino acids of SARS-2 FP1+, which include two additional non-polar residues (Val and Leu), generated only a small increase in the strength of the hydrophobic interaction (1.91 ± 0.03 to 2.21 ± 0.02 nN, Figure 3a). This small change in the hydrophobic interaction contrasts to the threefold effect of the Phe2Ala substitution in the FP1 sequence on the hydrophobic interaction strength.

**Figure 3.**
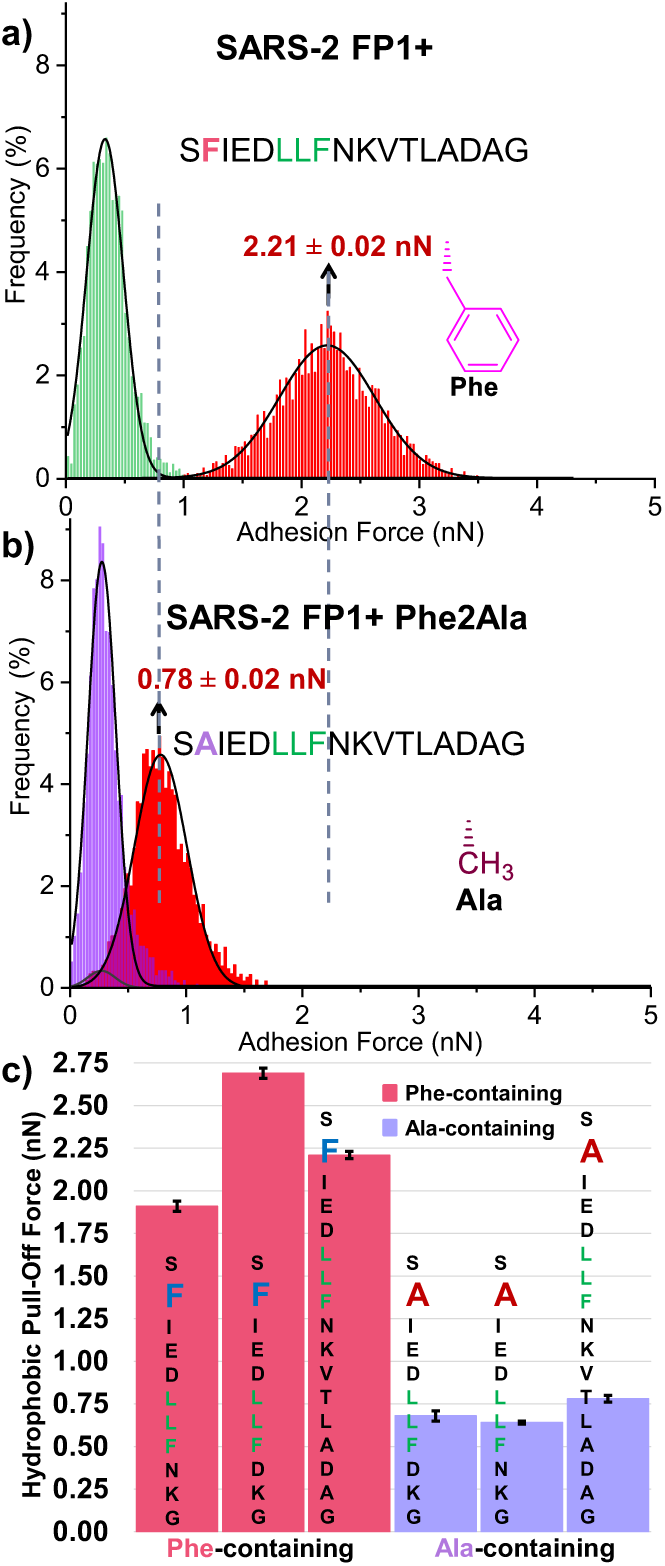
Pull-off forces measured using immobilized fusion peptides. a) Adhesive force histograms of SARS-CoV-2 fusion peptide (peptide sequence above) measured in PBS (red) and 60 volume % MeOH (green). b) Adhesive force histograms of SARS-CoV-2 fusion peptide with point mutation (peptide sequence Phe mutated to Ala) measured in PBS (red) and 60 volume % MeOH (purple). Dashed lines drawn for measurements in PBS to guide the eye. c) Comparison of adhesive forces of fusion peptide sequences containing Phe (pink bars) vs. Ala (purple bars). Adhesion force histograms and bar graphs were obtained using over 3,000 pull-off force curves from 6 independent samples. Data show mean ± s.e.m.

We next measured the effects of the Phe 2 to Ala 2 mutation on the hydrophobic interaction encoded by the SARS-2 FP1+. The strength of hydrophobic interaction encoded by the SARS-2 FP1+ Phe2Ala was measured to be 0.78 ± 0.02 nN (Figure 3b, red histogram), whereas the mean pull-off forces measured in the presence of 60 vol % MeOH decreased to 0.28 ± 0.01 nN (Figure 3b, purple histogram). In contrast, as reported above, the wild-type sequence with Phe 2 exhibits a threefold stronger hydrophobic interaction (2.21 ± 0.02 nN). The key conclusion emerging from this experiment is that the outsized influence of Phe 2 on the hydrophobic interaction mediated by the FP1 of SARS-CoV-2 is not limited to the FP1 sequence but is also observed with the longer sequence of 17 amino acids that contains a portion of the FP2. This point is shown in Figure 3c, which compares the hydrophobic interactions encoded by all six peptide sequences described so far in this paper. Significantly, all sequences that contain Phe 2 flanking LLF encode strong hydrophobic interactions; 1.91 ± 0.03 (SARS-2 FP1), 2.69 ± 0.03 (MERS FP1 Ala2Phe), and 2.21 ± 0.02 nN (SARS-2 FP1+) (Figure 3c, pink bars). In contrast, all sequences containing Ala 2 flanking LLF encode substantially weaker hydrophobic forces; 0.68 ± 0.03 (MERS FP1), 0.64 ± 0.02 (SARS-2 FP1 Phe2Ala), 0.78 ± 0.02 nN (SARS-2 FP1+ Phe2Ala) (Figure 3c, purple bars). Furthermore, a comparison of the hydrophobic interactions of the SARS-2 FP1+ Phe2Ala (0.78 ± 0.02 nN) and those of SARS-2 FP1 Phe2Ala which has the same FP1 amino acid composition (0.64 ± 0.02 nN) indicates that the additional non-polar residues of Ala-mutated SARS-2 FP1+ do not have a large impact the strength of hydrophobic interaction encoded by the FP sequences.

### Is LLF Important in Encoding the Hydrophobic Interactions of the FPs?

The results reported above also led us to explore the importance of LLF in determining the hydrophobic interactions encoded by the six amino acid sequences that we characterized. Past studies have proposed that LLF plays a critical role in determining viral membrane fusion (25,27,29,30). To evaluate the importance of LLF in encoding hydrophobic interactions, we performed a series of adhesion force measurements in which we replaced each of the three amino acids within the LLF triad with more polar (less non-polar) residues. For these experiments, we used the 17-amino acid FP sequence, as described above, with the SARS-2 FP1+ sequence serving as the reference (hydrophobic interaction of 2.21 ± 0.02 nN, Figure 4a, red histogram). The first mutation, which replaced Phe 8 of LLF with Tyr 8, resulted in a decrease in the strength of the hydrophobic interaction to 1.62 ± 0.01 nN (Figure 4b, red histogram). The second mutation involved the replacement of Leu 6 and Leu 7 with Ala 6 and Ala 7. This change resulted in hydrophobic interactions of strength 1.37 ± 0.01 nN (Figure 4c, red histogram). Finally, Leu 6 and 7 were replaced by Ser 6 and 7, resulting in hydrophobic interactions of 0.81 ± 0.01 nN (Figure 4c, red histogram).

Overall, these results reveal that substitution of LLF for amino acids that are more polar (less non-polar) incrementally weakened the hydrophobic interactions encoded by the FP sequence. The results thus confirm that LLF does play a key role in encoding the hydrophobic interaction of the FP1 sequence in our measurements, consistent with its reported role in studies of viral membrane fusion (25,27,29,30). These findings, when combined with the results shown in Figure 3, also hint that the hydrophobic interactions of FP1 from SARS-CoV-2 arise from a cooperative effect involving Phe 2 and LLF within the sequence (Figure 3c, pink bars).

### How does the FP1 Sequence Encode Hydrophobic Interactions?

To explore the physical mechanism by which Phe2Ala regulates the hydrophobic interaction encoded by LLF within the FP1 sequence of SARS-CoV-2 and MERS-CoV, we evaluated the hypothesis that Phe 2 exerts its influence via changes in the secondary structure of the oligopeptides. This hypothesis is based on the proposal that the conformations adopted by FPs while mediating hydrophobic interactions with an interface influence the nanoscopic patterns of non-polar amino acids presented at the interface.

The conformational states adopted by oliopeptides and proteins at interfaces can be characterized ATR-FTIR (56-61). In particular, the Amide I spectroscopic region (1600 – 1700 cm^-1^), which arises from stretching vibrations of peptide carbonyl groups, is sensitive to the peptide conformational state (62-64). Past studies have determined that peak positions in the Amide 1 region indicative of a β-sheet conformation are centered at 1624 cm^-1^; random coil at 1645 cm^-1^; α-helix at 1656 cm^-1^; and turns at 1670 cm^-1^ and 1680 cm^-1^ (62,65-68). We performed ATR-FTIR measurements using non-polar surfaces identical to the non-polar AFM tip surfaces. Briefly, we deposited a thin layer of gold onto a germanium ATR crystal, followed by adsorption of 1-dodecanethiol to form a non-polar monolayer. Finally, FP sequences were adsorbed onto the non-polar monolayer from PBS and ATR-FTIR measurements were conducted in PBS (Figure 5a). To avoid disulfide bond formation between thiol-capped peptides (as used in the AFM experiments above), the FP sequences used in ATR-FTIR measurements were capped with an acetyl group at the N-terminus. Below, we present ATR-FTIR spectra that show the Amide I peak region, with spectra obtained over a wider range of wave numbers presented in Supporting Information (Figures S3 and S4).

First, we used ATR-FTIR measurements to characterize the secondary structures of the adsorbed FP1 sequences that were used in the AFM measurements reported in Figure 2. Inspection of Figure 5b reveals an Amide I peak centered at 1632 ± 1 cm^-1^ for adsorbed MERS FP1 with shoulders at 1648 ± 1 cm^-1^, 1678 ± 1 cm^-1^, and 1717 ± 1 cm^-1^ (Figure 5b, pink curve; Table 1). This result suggests that the conformation of adsorbed MERS FP1 is dominated by a random coil state but also includes turns (as indicated by the shoulders). In contrast, the spectra obtained using the SARS-2 FP1 is clearly different, with an Amide I peak position at 1655 ± 1 cm^-1^, indicating α-helical content in the adsorbed state (Figure 5b, green curve; Table 1). This initial result provides support for our hypothesis that the SARS-2 FP1 and MERS FP1 sequences interact with non-polar surfaces via distinct conformational states.

**Table 1.** Amide I peak positions (cm^-1^) and corresponding hydrophobic pull-off forces (nN).

| | Wavenumber in PBS ($\text{cm}^{-1}$) | Adhesive Force in PBS (nN) | Wavenumber of Shoulder in PBS ($\text{cm}^{-1}$) |
| --- | --- | --- | --- |
| MERS FP1 | $1632 \pm 1$ | $0.68 \pm 0.03$ | $1648 \pm 2$;<br>$1678 \pm 1$;<br>$1717 \pm 1$ |
| SARS-2 FP1 | $1655 \pm 1$ | $1.91 \pm 0.03$ | |
| MERS FP1 Ala2Phe | $1660 \pm 1$ | $2.69 \pm 0.03$ | |
| SARS-2 FP1 Phe2Ala | $1639 \pm 1$ | $0.64 \pm 0.02$ | |
| SARS-2 FP1+ | $1657 \pm 1$ | $2.21 \pm 0.01$ | |
| SARS-2 FP1+ Phe2Ala | $1640 \pm 1$ | $0.78 \pm 0.02$ | |
| SARS-2 FP1+ Phe8Tyr | $1652 \pm 2$ | $1.62 \pm 0.01$ | |
| SARS-2 FP1+ Leu6Ala | $1644 \pm 1$ | $1.37 \pm 0.02$ | |
| SARS-2 FP1+ Leu6Ser | $1637 \pm 1$ | $0.81 \pm 0.01$ | $1678 \pm 1$ |

Next, we explored the effect of replacement of Ala 2 by Phe 2 on the conformational states of the adsorbed FP1 peptides. Inspection of Figure 5b reveals that MERS FP1 Ala2Phe exhibited a peak absorbance at 1660 ± 1 cm^-1^ (Figure 5b, blue curve; Table 1). This peak position is consistent with an α-helix, with the position of the peak shifted towards higher wavenumbers as compared to SARS-2 FP1 at 1655 ± 1 cm^-1^. This result indicates that while the peptide assumes an α-helix, turns are also present within the adsorbed peptide population on the non-polar interface. Finally, SARS-2 FP1 Phe2Ala exhibited a spectrum with a primary Amide I peak position at 1639 ± 1 cm^-1^ (Figure 5b, purple curve; Table 1), indicating a largely random coil conformation. Overall, this set of findings reveals that a single point mutation from Ala to Phe at position 2 exerts a pronounced influence over the FP1 conformation when mediating hydrophobic interactions, driving the FP1 sequence to switch in the adsorbed state from a largely random coil conformation to a largely α-helical conformation.

We performed a third series of ATR-FTIR measurements using the 17-amino acid SARS-2 FP1+ sequences, including mutations to the LLF triad reported in the context of Figures 3 and 4. Upon adsorption of SARS-2 FP1+ onto the non-polar surface of the ATR crystal, the Amide I peak absorbance was measured at 1657 ± 1 cm^-1^ (Figure 5c, mauve curve; Table 1). This result is similar to the Amide I peak of SARS-2 FP1 (Figure 5b, green curve; Table 1), indicating a largely α-helical conformation. Additionally, each mutation of LLF to polar (less non-polar) amino acids (to Tyr, Ala, and Ser, respectively) was measured to incrementally shifted the Amide I peaks towards smaller wavenumbers, indicating a transition towards random coil conformational states (Figure 5c, blue, green, and purple curves; Table 1). We also we examined the Amide I peak positions of SARS-2 FP1+ Phe2Ala, in which Phe 2 was substituted for Ala 2 (Figure 5c, orange curve; Table 1). We measured an Amide I peak at 1640 ± 1 cm^-1^, revealing a predominantly random coil conformation. These results, when combined with conclusions from ATR-FTIR measurements of FP1 sequences in Figure 5b, establish that both Phe 2 and LLF are needed to induce α-helical conformations of FP sequences at the non-polar surface; the absence of either of these two features of the peptide results in a random coil conformation. This result also emphasizes the interplay between the hydrophobic interaction and conformation, a point that we return to below.

**Figure 4.**
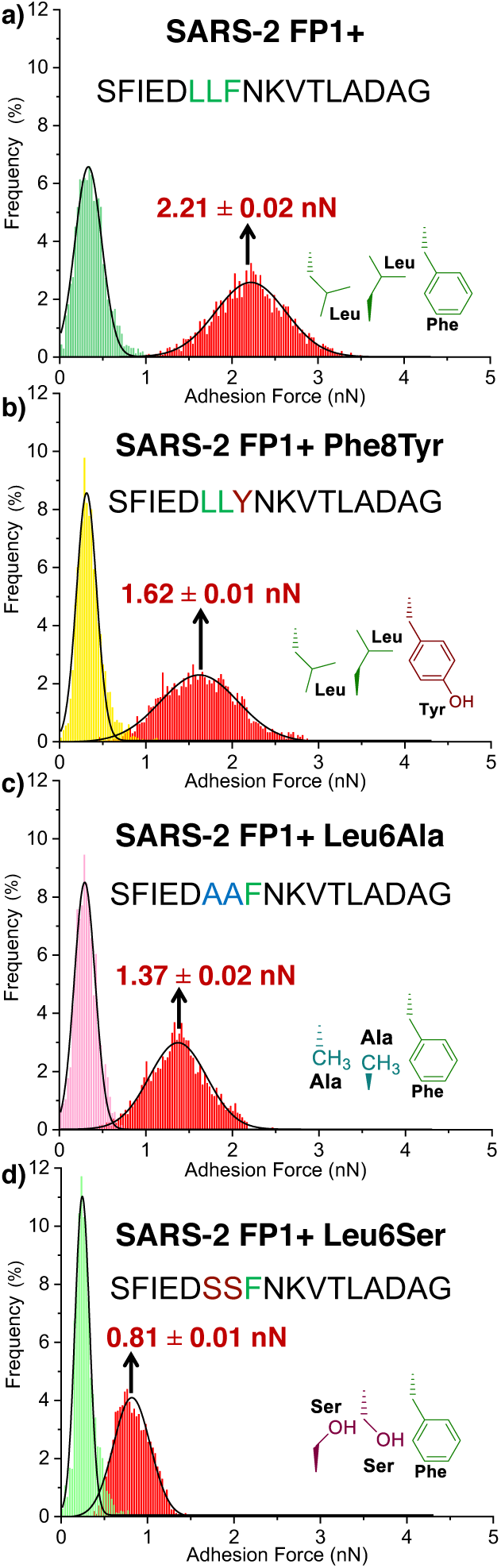
a) Peptide sequence from the wild-type FP of SARS-CoV-2 (S1 to G17) (top) and variants used to obtain the force histograms shown in (a-d). In the three sequences shown below the wildtype sequence in (a), we substituted LLF residues for the less non-polar amino acids of tyrosine (b), alanine (c), and serine (d). Adhesion force histograms were obtained using over 3,000 pull-off force curves from 6 independent samples. Data show mean ± s.e.m.

**Figure 5.**
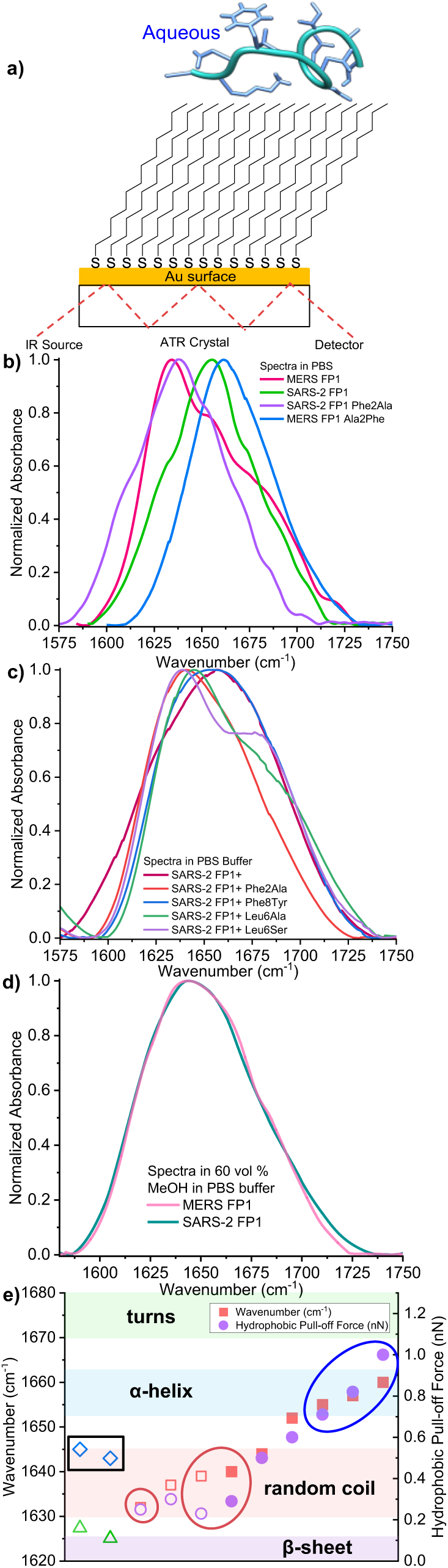
a) ATR-FTIR experimental setup. A thin layer of gold is electron beam deposited onto Ge ATR crystal, followed by adsorption of 1-dodecanethiol. Fusion peptides are then adsorbed onto the alkyl-terminated non-polar surface. b) Amide I peak spectra of the 11-amino acid FP1 sequences measured in PBS containing 0.9 mM Ca^2+^. c) Amide I peak spectra of 17-amino acid FP1-FP2 sequences with LLF substitutions. d) Amide I peak spectra of the 11-amino acid FP1 sequences measured in 60 vol % MeOH in PBS containing 0.9 mM Ca^2+^. e) Summary of hydrophobic pull-off force measured in PBS as a function of Amide I peak position wavenumber measured in PBS containing 0.9 mM Ca^2+^. Data points in the blue circle consist of FP sequences in which Phe 2 flanks LLF measured in PBS. Data points in the red circles represent FP sequences in which Ala 2 flanks LLF measured in PBS. Data points in the black box denote the wavenumber the Amide I peak of SARS-2 FP1 and MERS FP1 measured in 60 vol % MeOH in PBS containing 0.9 mM Ca^2+^. Error bars are included, but the size of the data points overlaps the size of the error bars. The spectrum of each FP consists of the average of three independently collected spectra. All ATR-FTIR curves are the average of three independently collected spectra.

Finally, we used FTIR measurement to explore the influence of the addition of methanol on the conformations of the adsorbed FP1 peptide sequences because past studies have reported that addition of methanol can promote stabilization of α-helices in bulk solution via a weakening of hydrogen bonding with the solvent (and an increase in intramolecular hydrogen bonding) (69-73). In particular, we wanted to determine if the conformational states of the FP1 peptide sequences in PBS differed from those measured in PBS containing 60 vol % MeOH. We measured the Amide I peak position of the MERS FP1 sequence adsorbed onto the non-polar monolayer from PBS containing 60 vol % MeOH to be located at 1643 ± 1 cm^-1^, indicative of a primarily random coil conformation (Figure 5d, pink curve; Table 2). This result reveals that addition of 60 vol % MeOH to PBS did not enhance the α-helical content of the adsorbed MERS FP1 relative to the conformation measured in PBS (peak at 1632 ± 1 cm^-1^, with shoulders 1648 ± 1 cm^-1^, 1678 ± 1 cm^-1^, and 1717 ± 1 cm^-1^) However, our measurements of the Amide I peak position for the adsorbed SARS-2 FP1 sequence in PBS to which 60 vol % MeOH was added, which was located at 1645 ± 1 cm^-1^, indicates the presence of a predominantly random coil structure upon adsorption onto the non-polar monolayer, in contrast to its predominantly α-helical conformation when adsorbed from PBS.

**Table 2.**
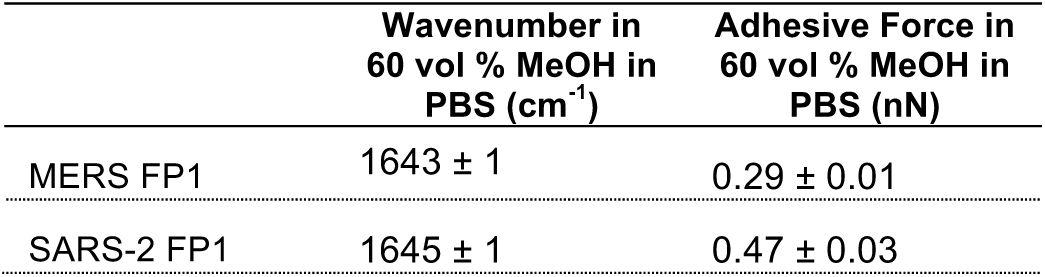
Peak positions of SARS-2 FP1 and MERS FP1 Amide I peaks (cm^-1^) and their corresponding adhesion forces (nN) in 60 vol % MeOH in PBS containing 0.9 mM Ca^2+^.

The results above, when combined, lead to two key observations. First, we observe the addition of 60 vol % MeOH to the PBS to promote the random coil conformational state of the adsorbed SARS-2 FP1 peptide sequence relative to the adsorbed conformational states measured in PBS alone. This result suggests that hydrophobic interactions do influence the conformations of adsorbed SARS-2 FP1 in PBS. It also contrasts to the previously reported effects of MeOH on the conformations of peptides in bulk solution (see below for measurements of CD spectra of the FP1 peptides in bulk solution). Second, in PBS, we observe a strong correlation between the secondary structure of adsorbed FP1 peptides and their hydrophobic interaction with the non-polar surface (Figure 5e). While similar correlations have been reported previously in contexts such as the interaction of antimicrobial peptides with non-polar surfaces (74,75), what is striking and distinct in our results in the dominant role of Phe 2 versus Ala 2 in determining both the conformation and hydrophobic interaction of the FP1 sequence with the non-polar surface (see SI for additional discussion).

As discussed above, our ATR-FTIR measurements performed with and without 60 vol % MeOH added to PBS suggest that the interaction of the SARS-2 FP1 sequence with the non-polar surface of the ATR-FTIR crystal plays a key role in determining the conformations of the adsorbed peptides. Here we consider these observations in light of past studies that have reported that amino acid residues within oligopeptides have a propensity to promote specific secondary structures in bulk solution (76-79). According to these prior studies, the amino acids involved in the single point mutations in our study are predicted to exhibit the following decreasing order of helical propensity (measured in kcal/mol): Ala (0), Leu (0.21), Ser (0.50), Tyr (0.53), Phe (0.54) (76). This ranking leads to the prediction that FP sequences containing Ala will adopt α-helical structures more readily than Phe-containing sequences in bulk solution, a prediction that does not correlate with our ATR-FTIR measurements of the surface-adsorbed FPs (MERS FP1 versus SARS-2 FP1).

To explore the conformations of the FP sequences in bulk solution, we performed CD spectroscopy in PBS. Prior studies have reported that α-helices exhibit negative spectroscopic bands at 208 nm and 222 nm, while random coils or disordered structures contain very low ellipticity above 210 nm (80). Figure 6 shows CD spectra of the FP sequences used in our study. In PBS at pH 7.4, both MERS FP1 (red spectrum) and SARS-2 FP1 Phe2Ala (green curve) generate spectra consistent with random coil conformations (Figure 6a). In contrast, the spectra of SARS-2 FP1 (blue spectrum) and MERS FP1 Ala2Phe (orange spectrum) are indicative of mixed random coil and α-helical content, as identified by the band at 208 nm and a weaker band at 222 nm. While the CD spectra of the latter two FP1 sequences do not indicate well-formed α-helical conformations, they do indicate a greater α-helical content than MERS FP1 and MERS FP1 Ala2Phe in bulk PBS.

**Figure 6.**
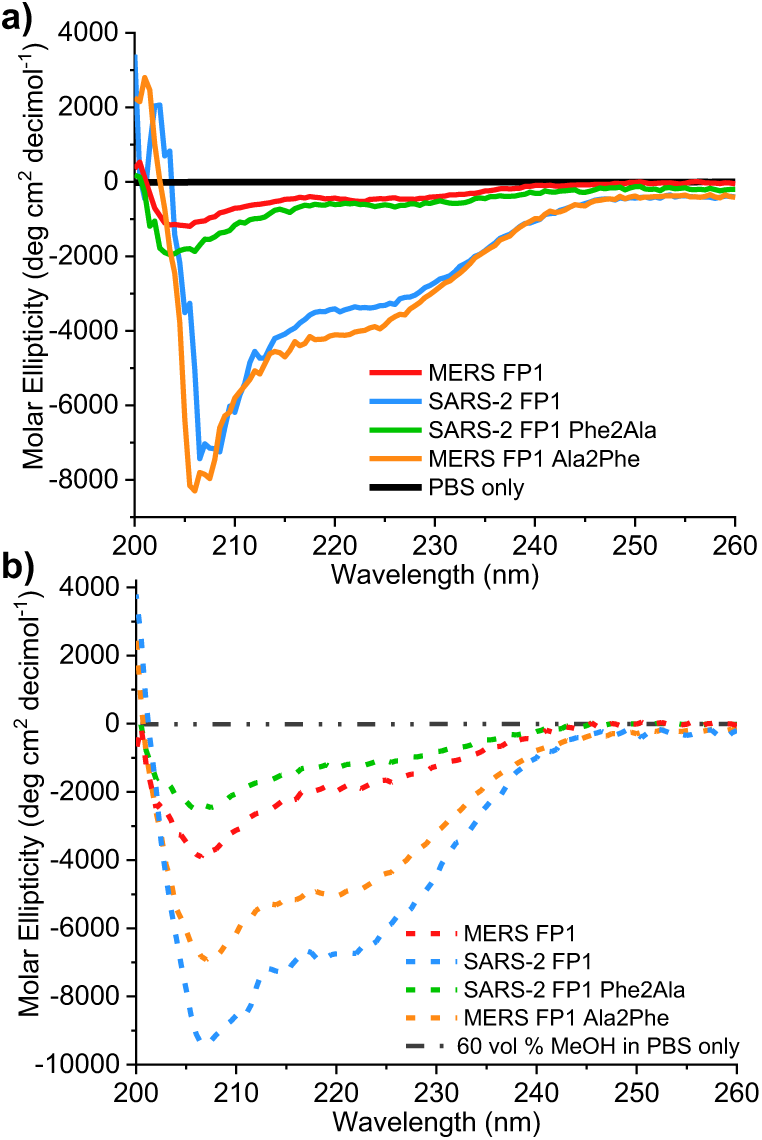
Circular dichroism of spectra of the 11-amino acid FP1 sequences in bulk PBS containing 0.9 mM Ca^2+^ (a) and 60 vol % methanol in PBS containing 0.9 mM Ca^2+^ (b). The spectra were normalized to mean residue ellipticity in units of deg cm^2^ decimol^-1^ (See Supporting Information Figure 8 for measurements in ellipticity). All CD curves are the average of three independently collected spectra.

From the measurements described above, in bulk PBS, we conclude that FP1 sequences with Phe 2 exhibit greater α-helical content than sequences with Ala 2. We estimated the percentage of α-helical content of sequences containing Phe 2 to be 8%, while that of sequences containing Ala 2 to be 1% (See Methods) (53,54). Thus, while the hydrophobic interactions of FP1 with surfaces in AFM and ATR-FTIR measurements play a role in determining the conformations of the peptides (as evidenced by the effect of adding 60 vol % MeOH on the conformation of the adsorbed SARS-2 FP1 sequence (Figure 5d, teal spectrum)), prior to contact with the surfaces, the peptides show weak preferences for distinct conformations. We also performed CD measurements on the 17-amino acid FP sequences introduced in Figure 4. Measurements revealed that SARS-2 FP1+ possesses weak α-helical character in bulk PBS, and that replacement of LLF in the SARS-2 FP1+ sequence led to a decrease in the α-helical content (see Supporting Information Figure S6a for further discussion).

We considered the possibility that the α-helical content of FP sequences in which Phe 2 flanks LLF may reflect hydrophobically-driven self-association of the FPs in bulk PBS. To address this possibility, we performed CD measurements in PBS with 60 vol % MeOH to probe conformational changes upon addition of methanol (Figure 6b; Supporting Information Figure 6b). Addition of methanol to aqueous buffer has been shown to disrupt hydrophobically-driven assembly (69-73). If the presence of α-helicity in FP sequences in our experiments is due to hydrophobically-driven association, addition of methanol would be predicted to disrupt the assembly and diminish the difference in CD spectra among sequences containing Ala 2 vs. Phe 2 in 60 vol % MeOH. However, we observed the differences between spectra obtained using sequences containing Ala 2 (dashed red and green) vs. Phe 2 (dashed blue and orange) in PBS to be maintained when 60 vol % MeOH was added to PBS (Figure 6b). in addition, we measured CD spectra of SARS-2 FP1 and MERS FP1 sequences in PBS at concentrations of 10, 100, and 1000 μM to evaluate if peptide self-association underlies the differences in CD signatures between sequences containing Phe 2 vs. Ala 2 in Figure 6a. We found no significant difference in CD signatures in spectra converted to mean residue ellipticity across the concentrations of each FP1 sequence (Supporting Information Figure 7c). We estimated the α-helical content of SARS-2 FP1 to be 8% at 10 and 100 μM, and 9% at 1000 μM (Supporting Information Figure 7b, solid curve), while that of MERS FP1 to be 1% at all three peptide concentrations (SI Figure 7b, dashed curves)(53,54). This result suggests that our CD measurements of Phe 2-containing sequences in PBS reflect the conformations of monomeric peptides, rather than self-associated complexes of peptides (see Supporting Information for further discussion).

## CONCLUSIONS

This study demonstrates that single-molecule force measurements permit quantification of the hydrophobic interactions encoded by FP sequences from SARS-CoV-2 and MERS-CoV. The measurements reveal that the non-polar triad Leu-Leu-Phe (LLF), which is conserved in both FP sequences, plays a central role in encoding the hydrophobic interaction of the FP sequences. This is consistent with prior studies that have concluded it to be a key determinant of membrane fusion between viral and host cell membranes (25,27,29,30). Surprisingly, however, we find that single amino acid residue differences within the FP1 sequences from SARS-2 and MERS, which are adjacent to LLF, can substantially alter the strength of the hydrophobic interaction mediated by the LLF. Specifically, we observe that the presence of Phe 2 in SARS-CoV-2 increased the magnitude of the hydrophobic interaction encoded by the FP by nearly a factor of 3 (in comparison to Ala 2 in MERS-CoV). Additionally, by performing ATR-FTIR measurements, we found strong support for the conclusion that Phe 2 exerts its outsized influence on the hydrophobic interaction encoded by LLF within the FP by regulating the secondary structure of the FP during hydrophobic interaction with surfaces. Specifically, the ATR-FTIR spectra of FP sequences with Phe 2 contained Amide I peaks at positions indicative of α-helical-rich conformational states, while FP sequences containing Ala 2 generated Amide 1 peaks at positions in the spectra indicative of largely random coil conformations. Our results reveal that single amino acid substitutions, i.e. switching between Phe 2 to Ala flanking LLF, can profoundly influence the secondary structure of peptides in the adsorbed state and the strength of hydrophobic interactions encoded by the FP.

The results of this study provide fresh ideas regarding factors that regulate hydrophobic interactions encoded by FPs of SARS-2 and MERS. The measurements, which were performed by interacting the peptides with non-polar surfaces to unmask their hydrophobic interactions, provide fundamental insights that can be used to design future studies of interactions of FPs with surfaces with the compositional complexity and dynamics characteristic of host cell membranes. In addition, we note that the experiments reported in this paper focused on FP sequence S1-S10 and S1-G17. Our results inform future studies of the full FP sequence consisting of 40 amino acids.

Hydrophobic interactions have been proposed to play a key role in driving membrane fusion between viruses and host cells (31-33). Accordingly, advances in our understanding of the mechanisms by which FP sequence impacts the interactions that drive fusion, such as those elucidated in this study, have the potential to inform strategies for designing molecules (other peptides or small molecule drugs) that modulate the interactions responsible for viral infection (5,12,81,82). In addition, our discovery of the impact of single amino acid substitutions on hydrophobic interactions encoded by the FP1 domain of CoVs provides new guidance to the judicious placement of residues in peptide sequences to modulate the conformations and interactions of peptides. These design rules have the potential to be useful not only for oligopeptides involved in viral fusion (83,84) but also for broader classes of peptide therapeutics and materials (85,86)

## SUPPORTING MATERIAL

Supporting material can be found online at doi.org/xxxxxx

## AUTHOR CONTRIBUTIONS

C.Q. performed AFM, ATR-FTIR, and CD measurements. C.Q., S.D., S.H.G. and N.L.A. designed research. C.Q, G.R.W., S.D., and N.L.A. analyzed data. C.Q. and N.L.A. wrote the manuscript. All authors participated in reviewing/editing of the manuscript and approved of the final draft.

## ACKNOWLEDGMENTS

This research was primarily supported by the NSF (MCB 2027070). The authors thank Miya Bidon for participating in helpful discussions. We also thank Mingxin He for his help in performing statistical analyses with *t*-tests.

