## Supporting Information file for "Characterization of Hydrophobic Interactions of SARS-CoV-2 and MERS-CoV Spike Protein Fusion Peptides Using Single Molecule Force Measurements"

| **The number of test events / The number of samples (n / N)** | | |
| --- | --- | --- |
| **Fusion Peptide** | **PBS** | **60 vol % MeOH in PBS** |
| MERS FP1 | 3240 / 6 | 3314 / 6 |
| SARS-2 FP1 | 3109 / 6 | 3261 / 6 |
| MERS FP1 Ala2Phe | 3822 / 6 | 3103 / 6 |
| SARS-2 FP1 Phe2Ala | 3517 / 6 | 3411 / 6 |
| SARS-2 FP1+ | 3221 / 6 | 3192 / 6 |
| SARS-2 FP1+ Phe2Ala | 3481 / 6 | 3321 / 6 |
| SARS-2 FP1+ Phe8Tyr | 3004 / 6 | 3163 / 6 |
| SARS-2 FP1+ Leu6Ala | 3190 / 6 | 3452 / 6 |
| SARS-2 FP1+ Leu6Ser | 3801 / 6 | 3617 / 6 |

**Table S1.** Statistical information for data shown in Figure 2-4, Table 1, Table 2. Total number of force measurements (n) and number of independent samples (N) for each fusion peptide interaction with AFM tip.

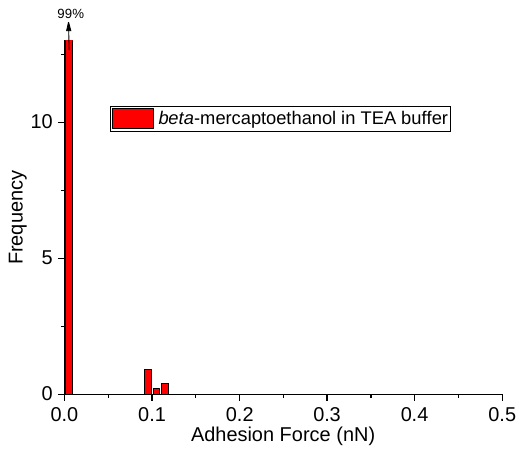

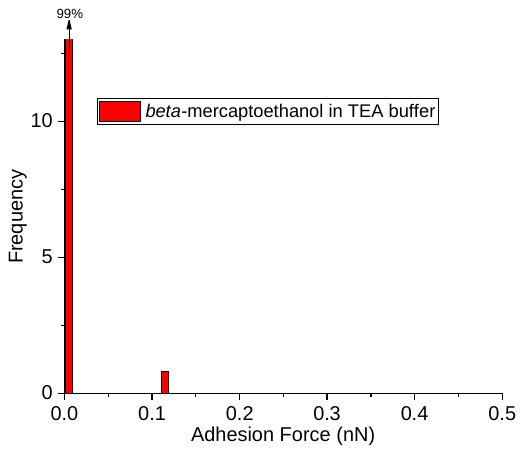

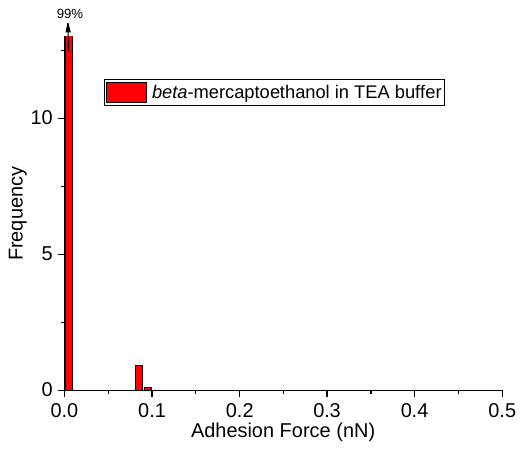

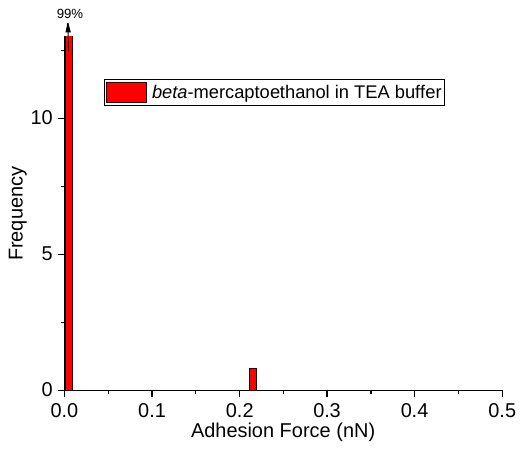

**Figure S1.** Adhesion force histograms of interactions between AFM tips chemically modified with adsorption of 1-docecanethiol and surfaces covalently immobilized with β-mercaptoethanol. Surfaces were activated with SSMCC, followed by covalent attachment of β-mercaptoethanol. Finally, a solution of thiol-terminted FP was incubated against the surface containing β-mercaptoethanol. The thiol group of β-mercaptoethanol should react with the maleimide groups of the SSMCC units, thereby preventing covalent attachment of FPs after pretreatment. Pretreatment with β-mercaptoethanol gave rise to largely non-adhesive interactions, shown in above histograms. Measurements were performed in aqueous PBS buffer at pH 7.4 for all four independently measured events. Adhesion forces histograms were obtained using over 3,000 pull-off force curves for each of the four independent samples.

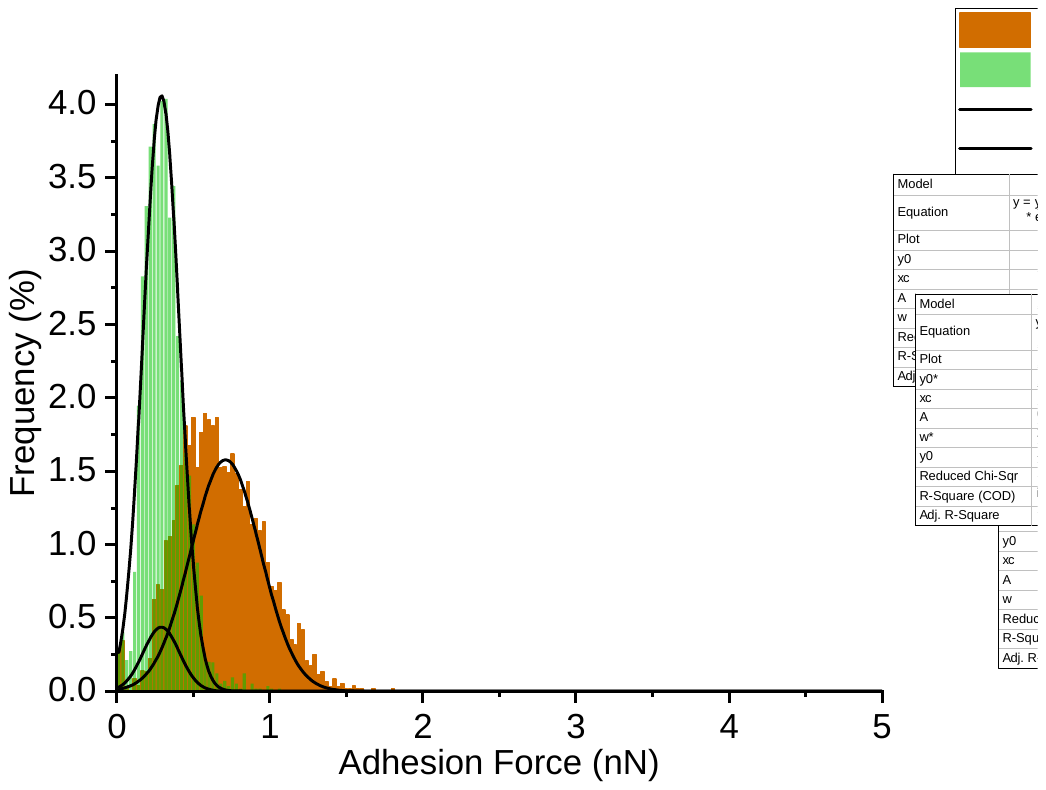

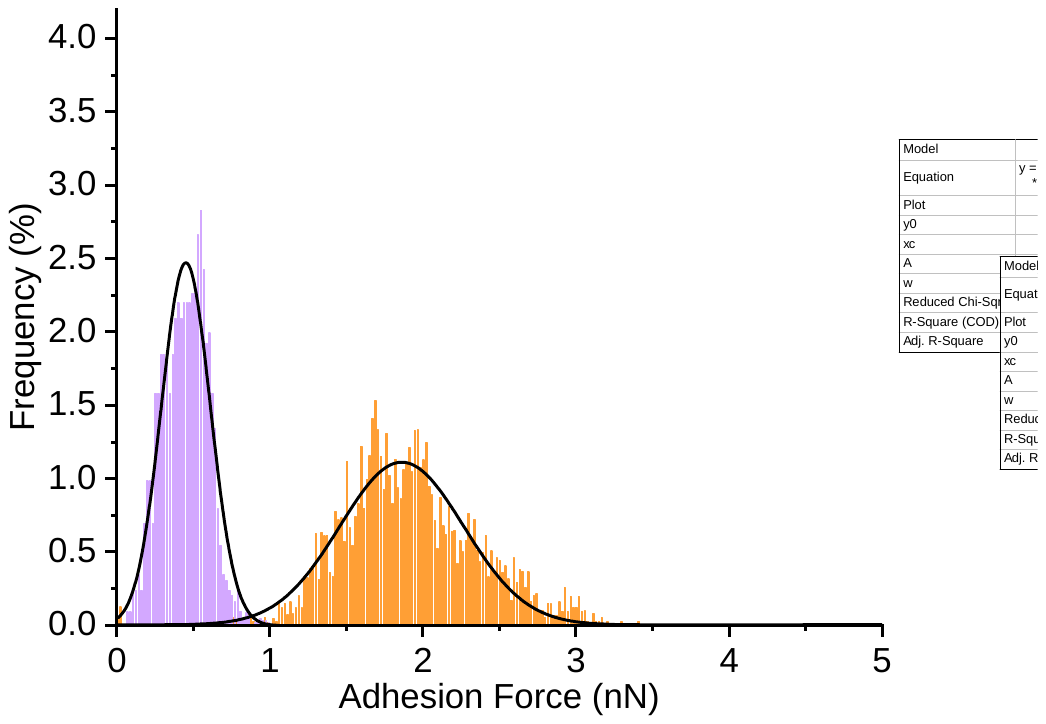

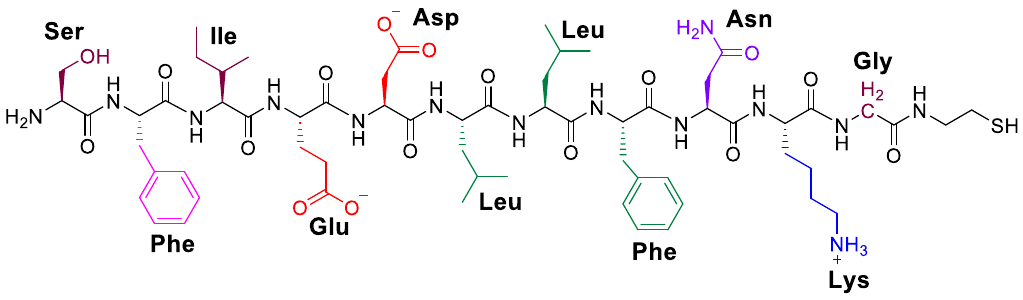

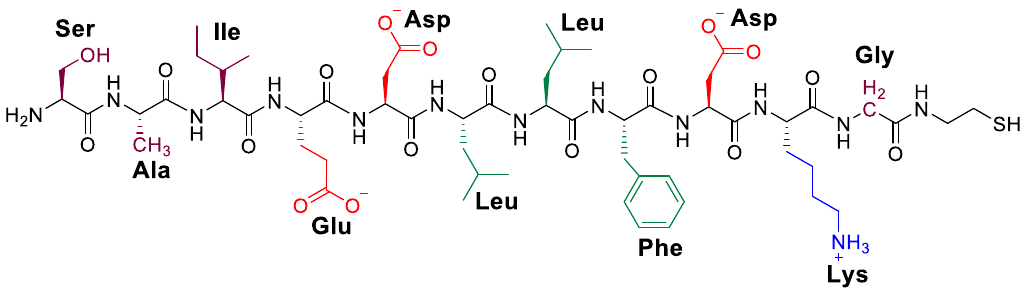

b)

a)

**Figure S2.** Adhesion force histograms of interactions between a nonpolar AFM tip and SARS-2 FP1 (a) and MERS FP1 (b) immobilized at a density of 0.001 mole fraction of aminotetraethylene glycol (EG4N) groups, in contrast to a mole fraction of 0.002 EG4N peptide immobilization density in Figures 2-4 in the main text. Adhesion force of SARS-2 FP1 and MERS FP1 in aqueous PBS buffer containing 0.9 mM Ca^2+^ (orange and brown histograms for SARS FP1 and MERS FP1, respectively). The mean adhesion force in PBS of SARS-2 FP1 is 1.86 ± 0.02 nN and that of MERS FP1 is 0.71 ± 0.01 nN.The purple (SARS-2 FP1) and green (MERS FP1) histograms show adhesion forces upon addition of 60 vol % MeOH in PBS. The mean adhesion force in PBS containing 60 vol % MeOH of SARS-2 FP1 is 0.45 ± 0.02 nN and that of MERS FP1 is 0.29 ± 0.02 nN. Adhesion force histograms were obtained using over 3,000 pull-off force curves from 6 independent samples.

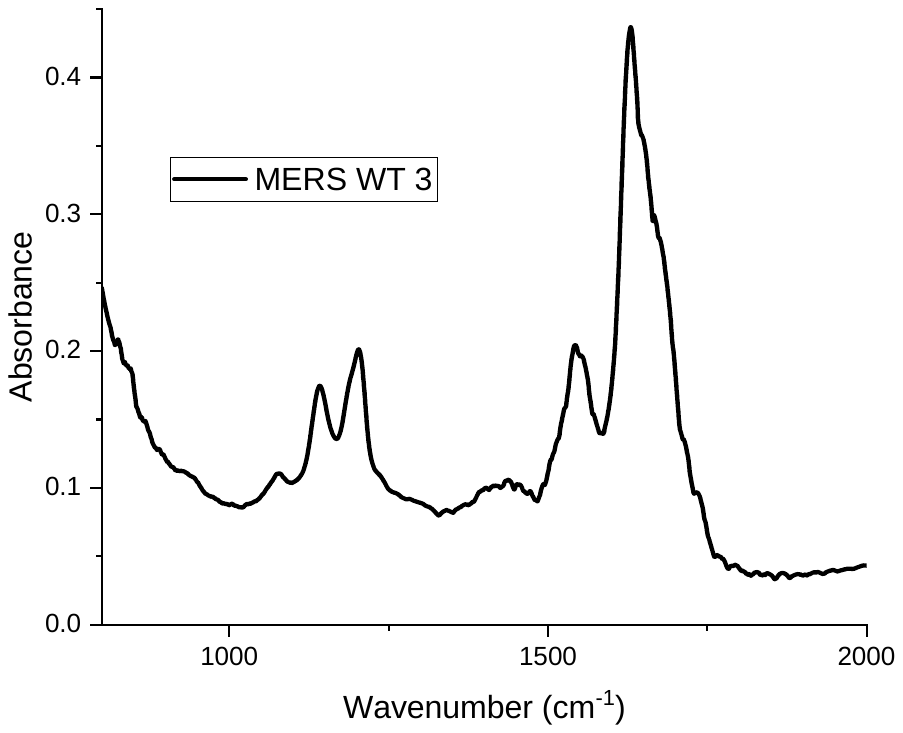

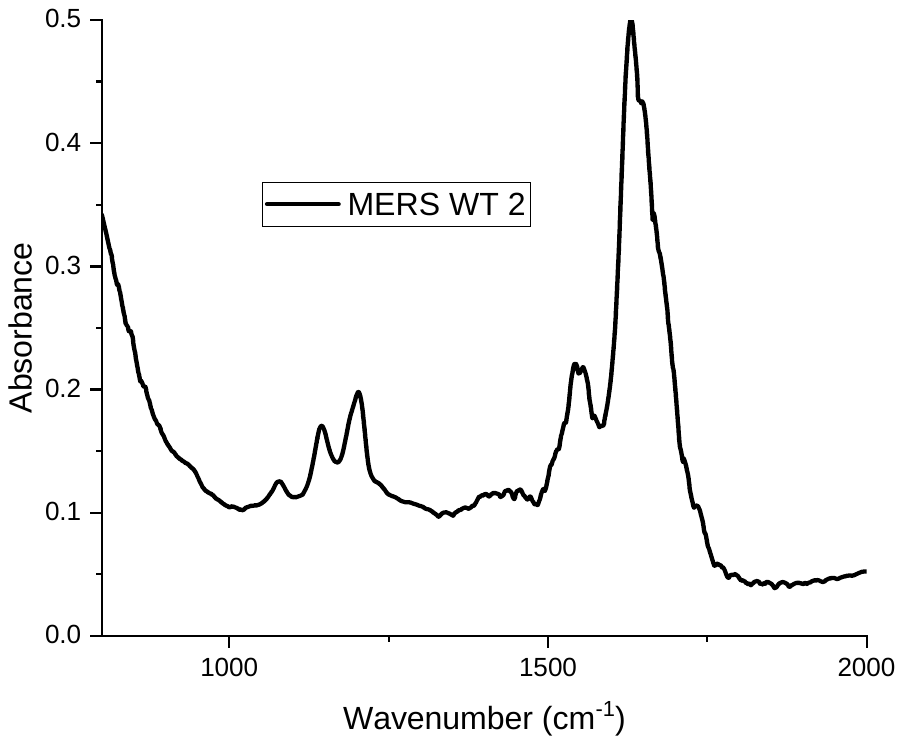

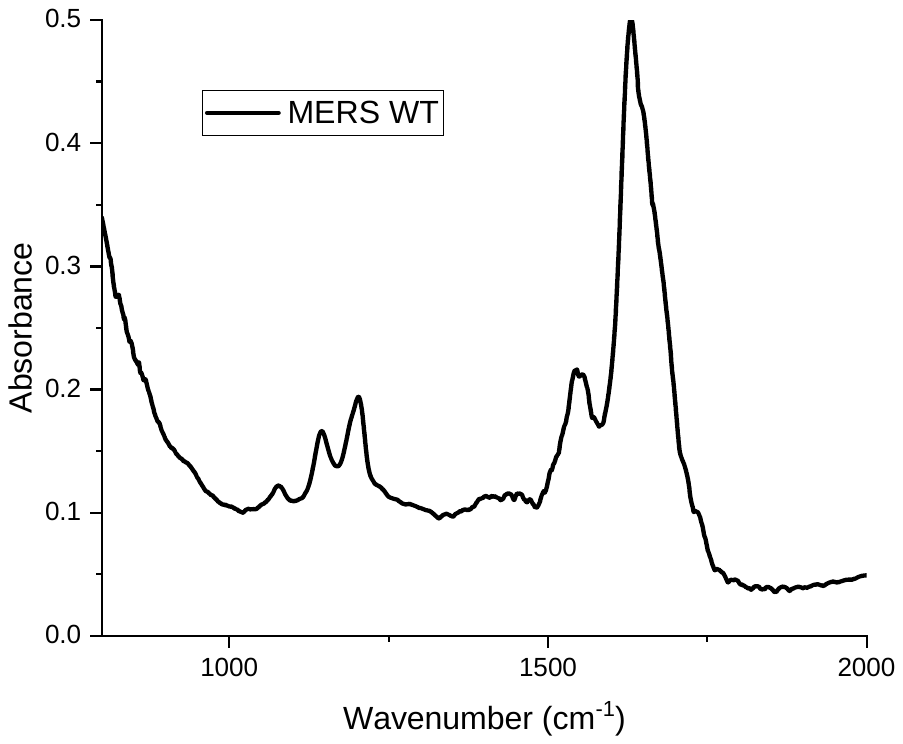
a)

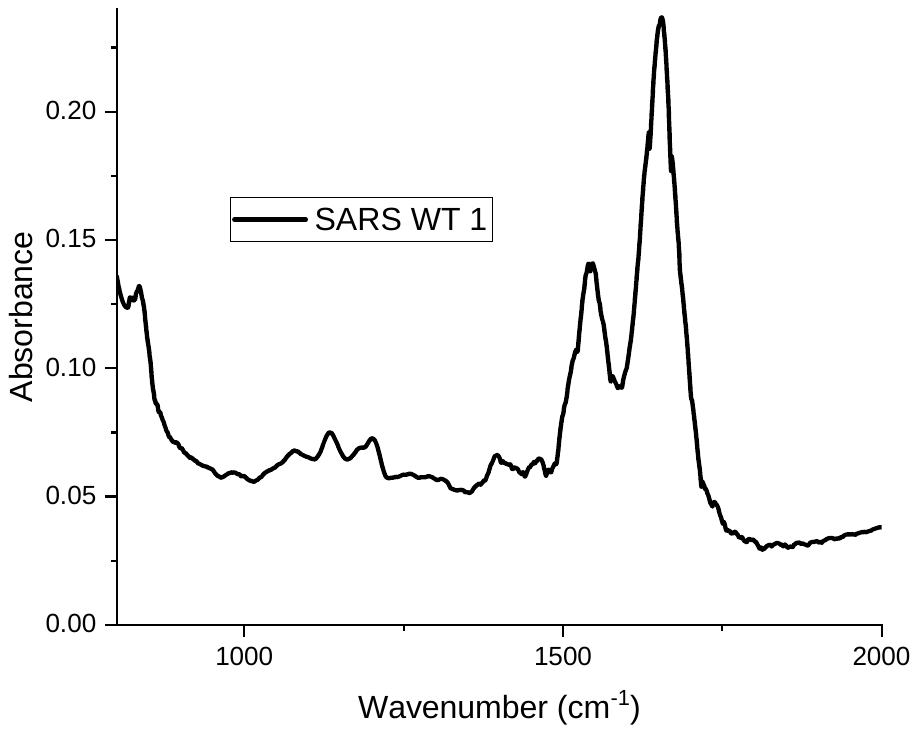

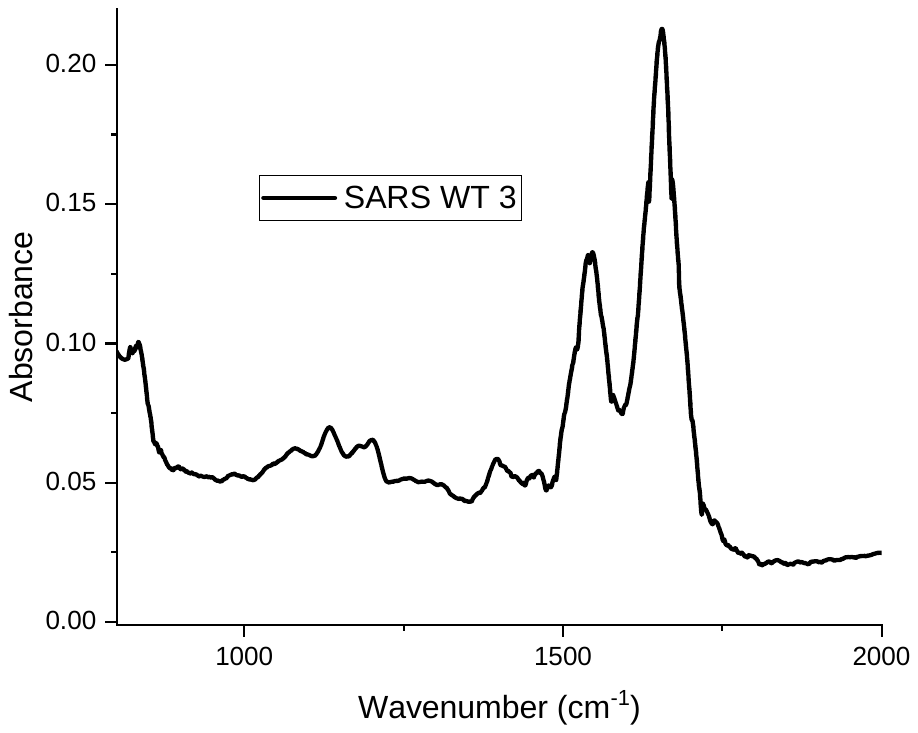

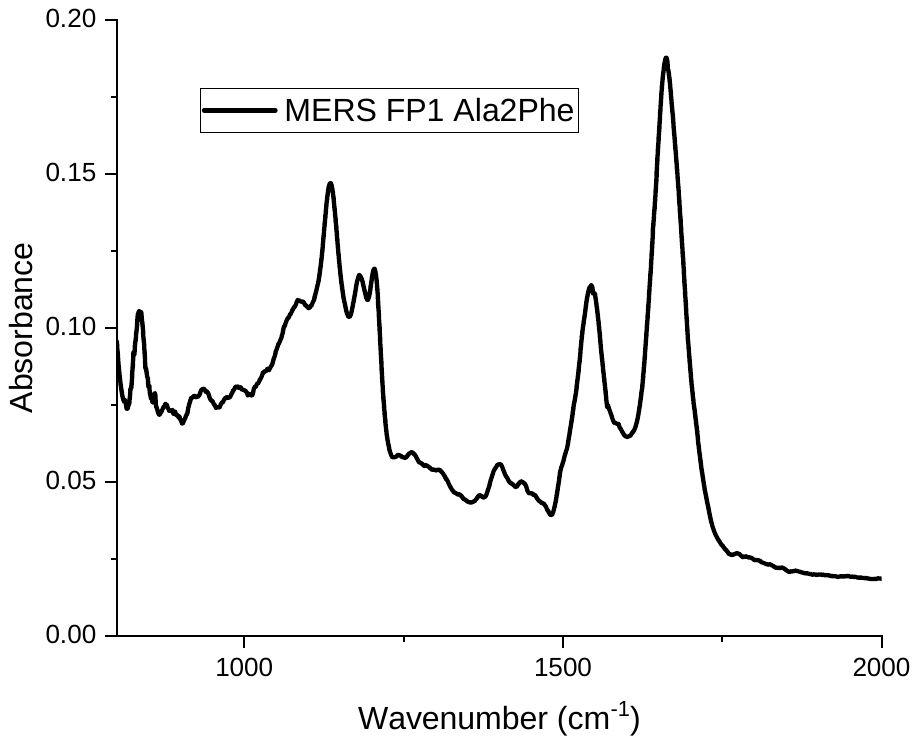

b)

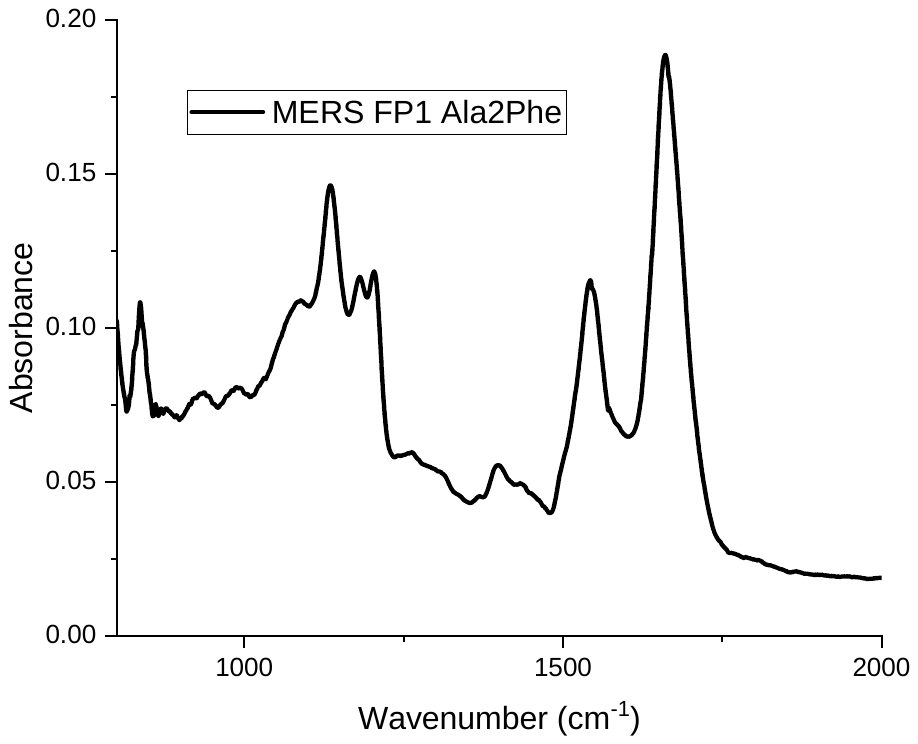

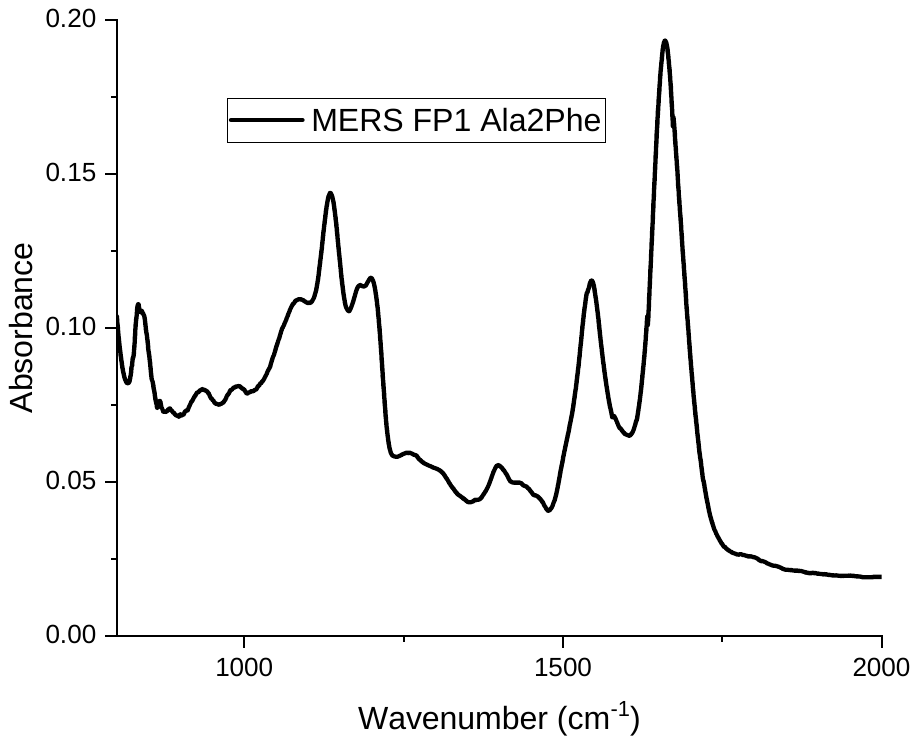

c)

**Figure S3.** Extended ATR-FTIR spectra measured in PBS containing 0.9 mM Ca^2+^ of 11-amino acid FP1 sequences of SARS-2 FP1 wild-type (a); MERS FP1 wild-type (b); MERS FP1 Ala2Phe (c). The peptide backbone give rise to amide I (1600 – 1700 cm^-1^), corresponding to C=O stretching; amide II (1480 – 1575 cm^-1^), corresponding to N-H bending and C-N stretching; and amide III (1200 – 1350 cm^-1^), corresponding to C-N stretching, N-H bending, C-C stretching, and C-H bending. Of particular relevancy to peptide secondary structure are amide I and amide II bands. Since both C=O and N-H bonds are involved in hydrogen bonding, which gives insight into the secondary structure, amide I and amide II bands are generally used to yield information about peptide conformation (1,2). Among these amide bands, amide I is most widely used to determine peptide secondary structure as a result of its strong signal. Three individual spectra were collected for each FP and averaged to yield the spectra shown in Figure 5c of the main text.

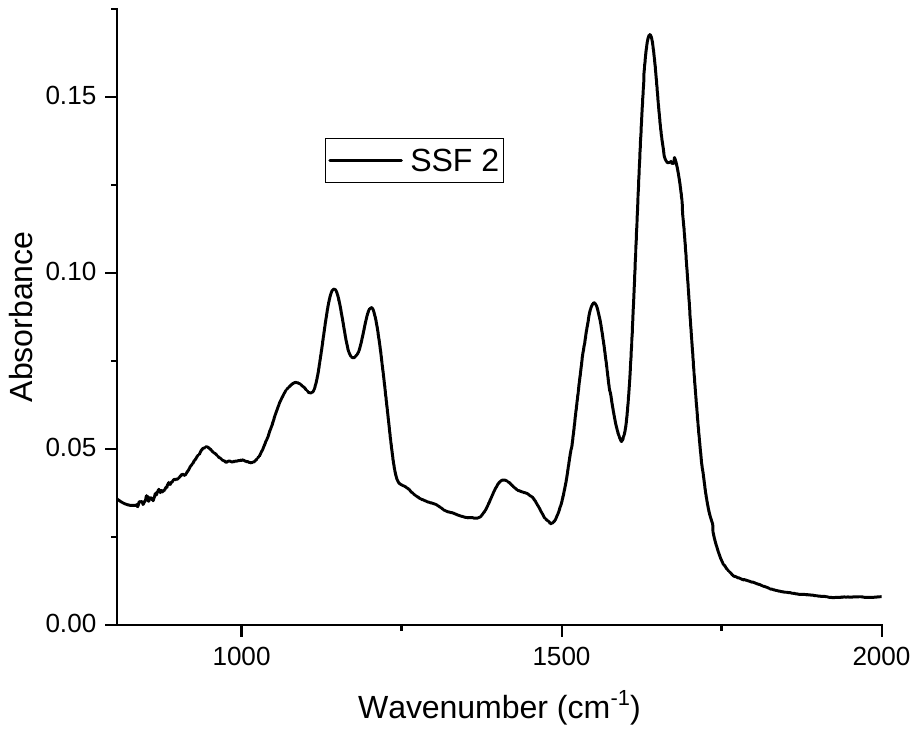

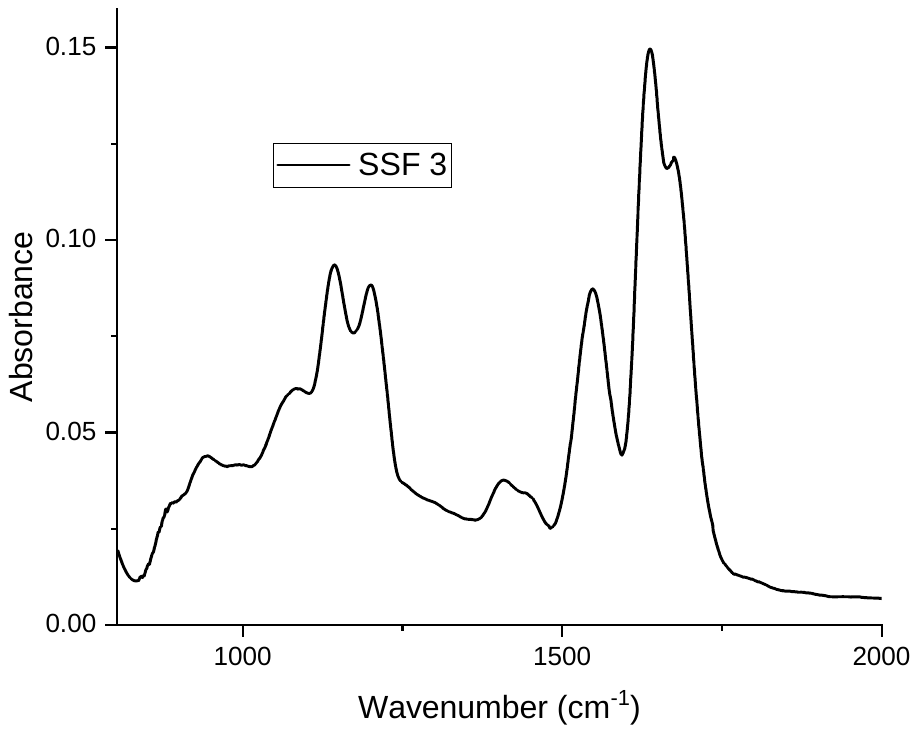

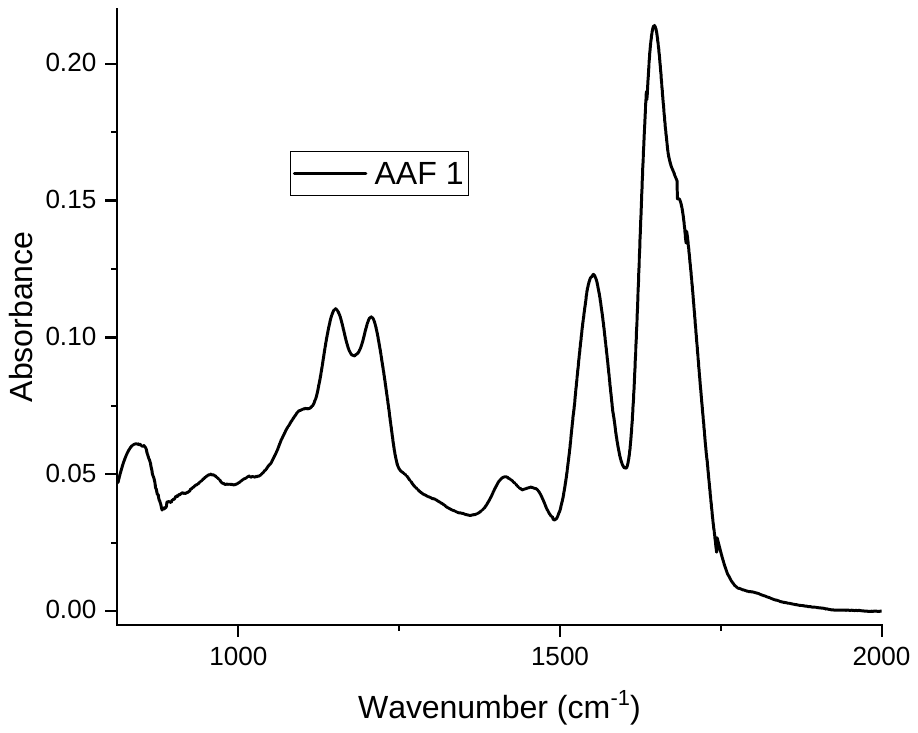

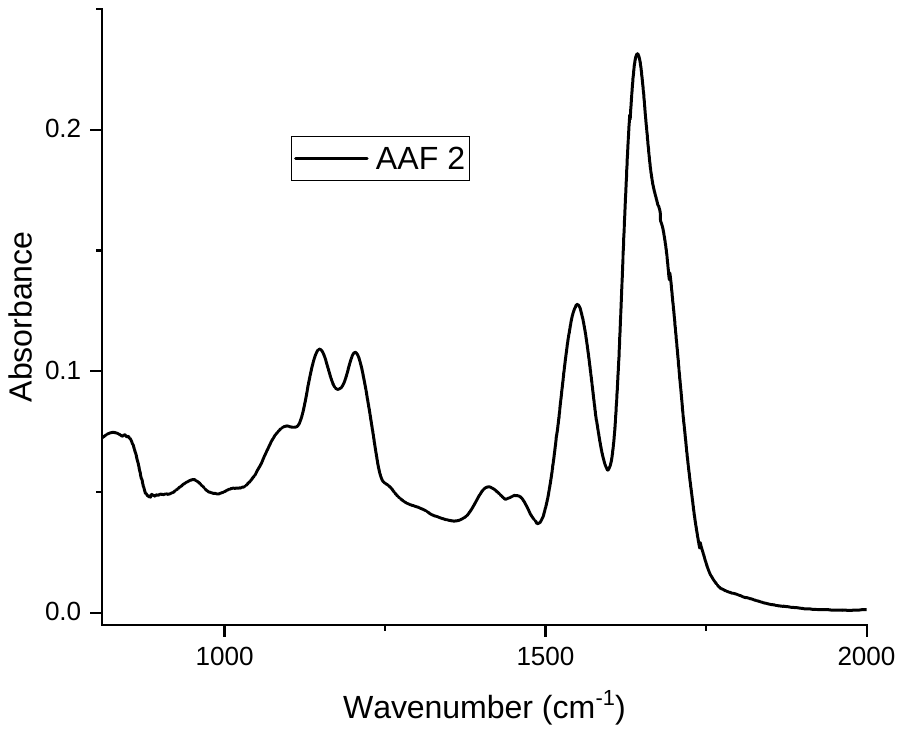

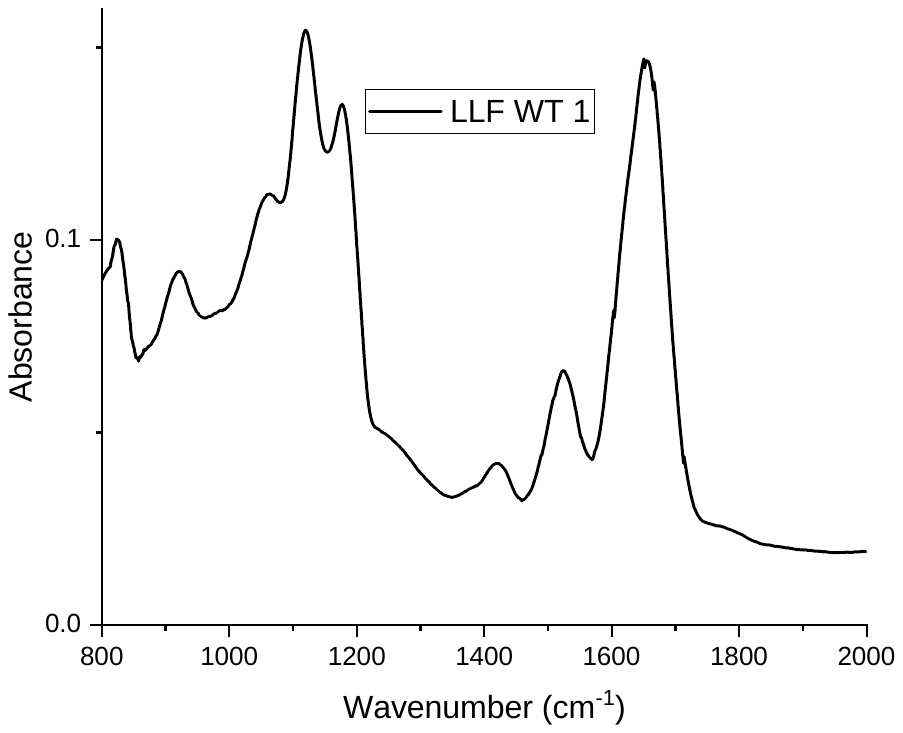

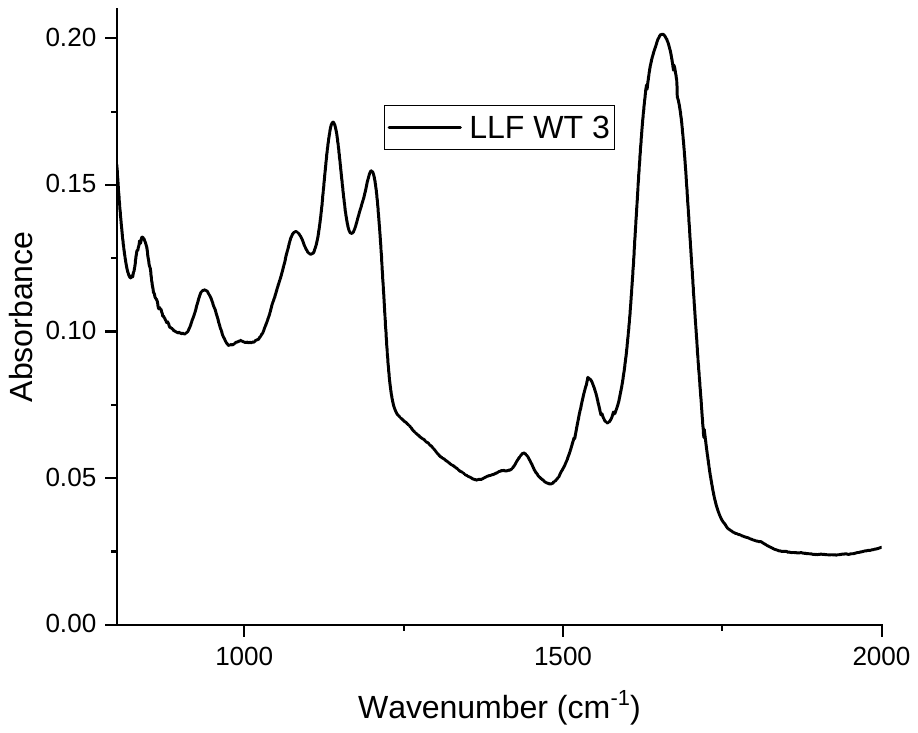

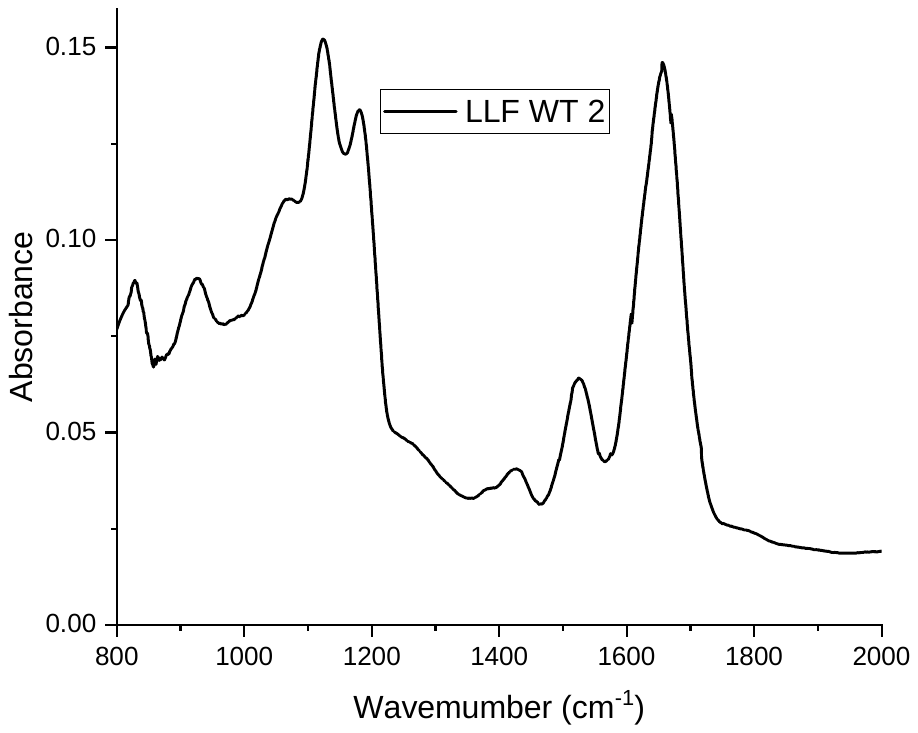

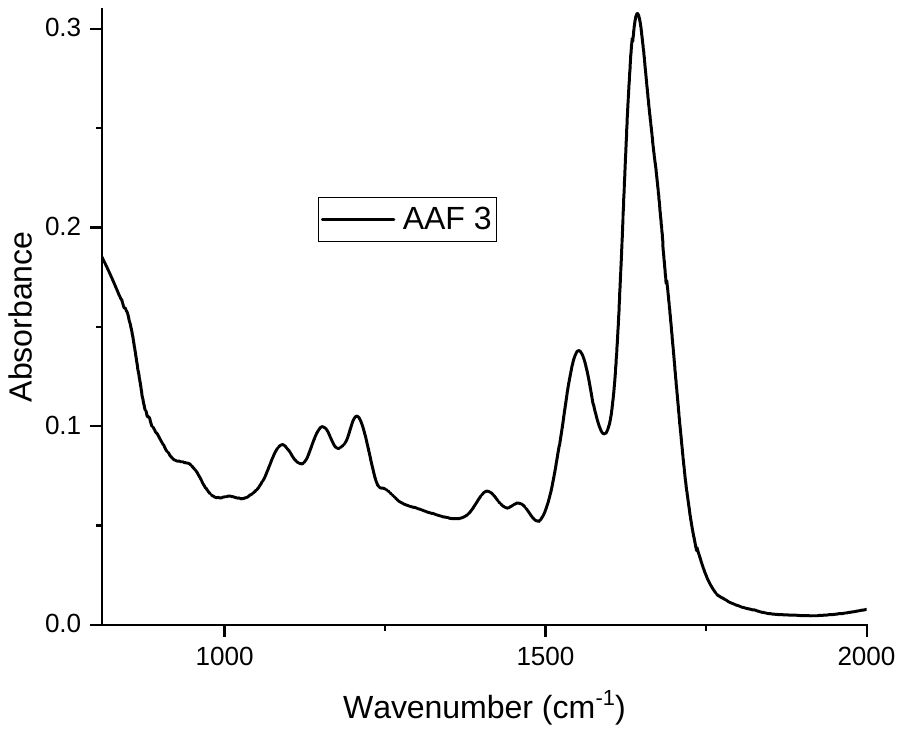

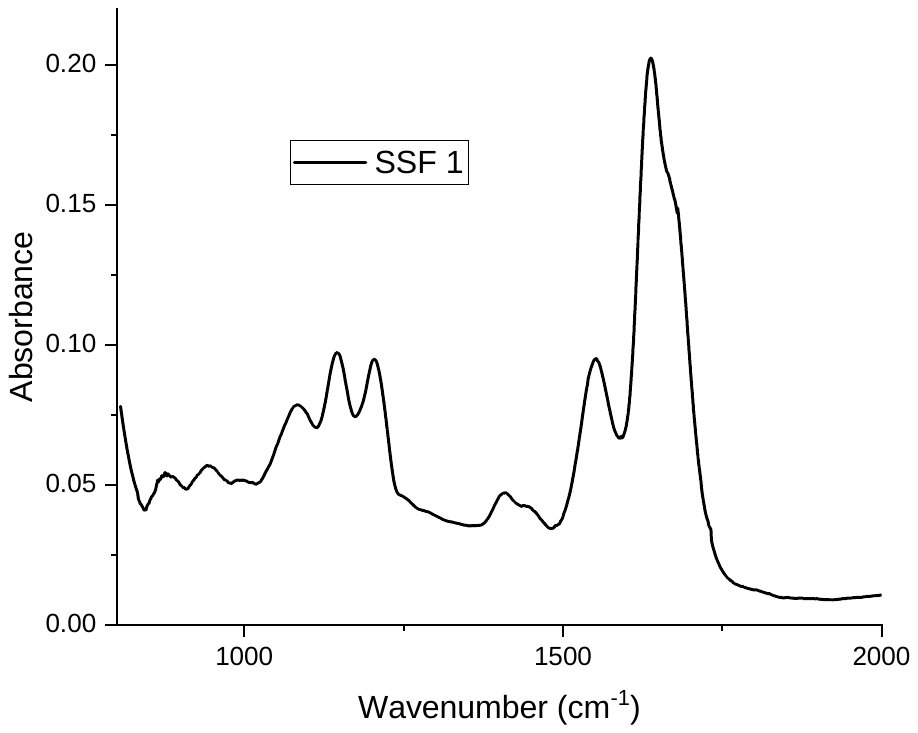

a)

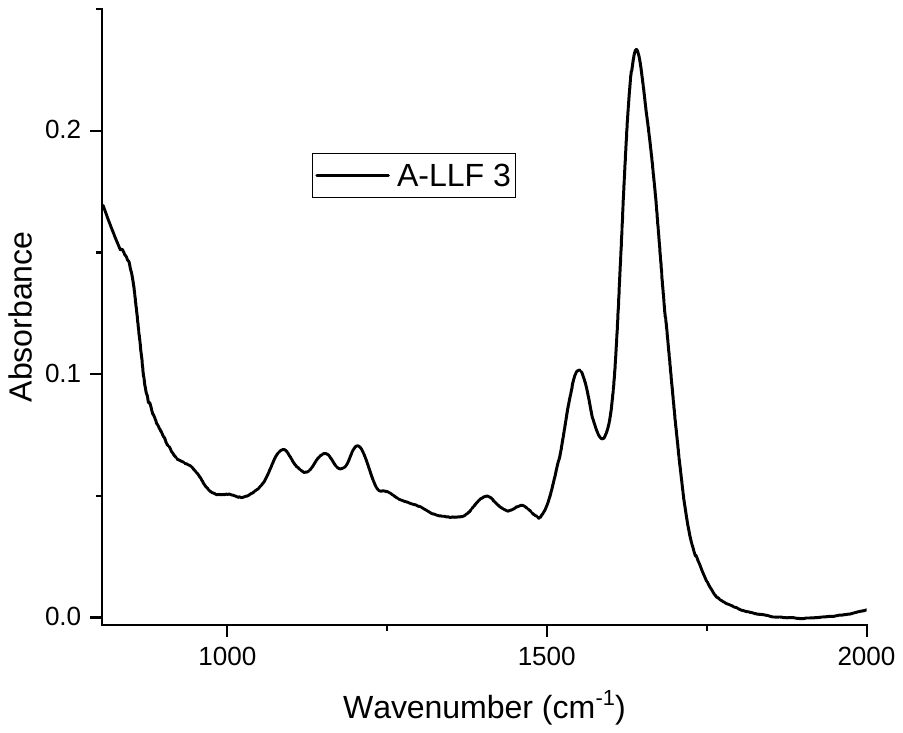

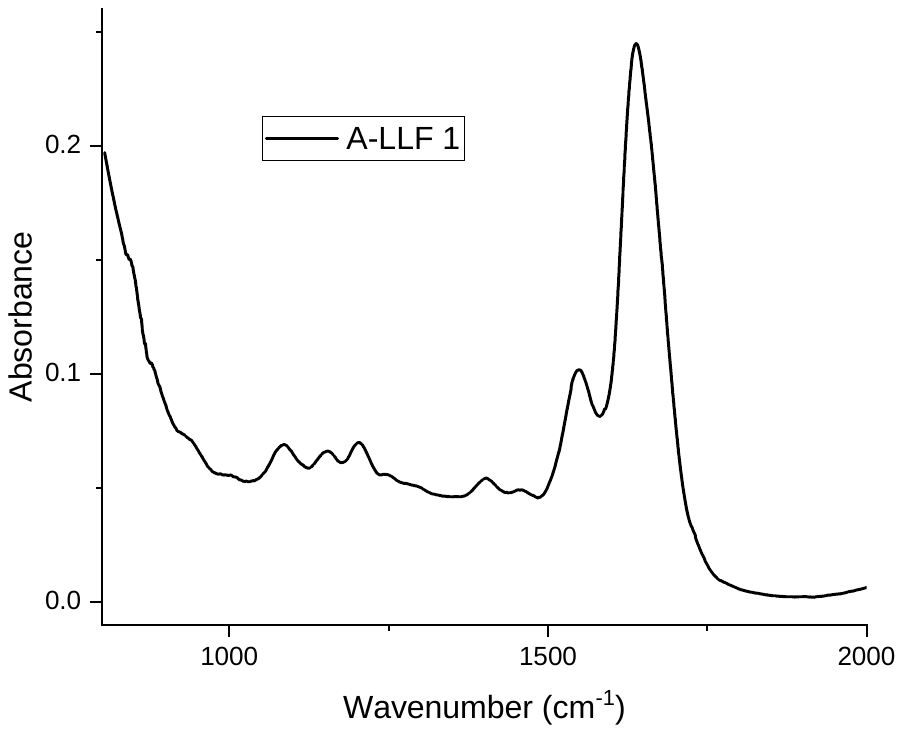

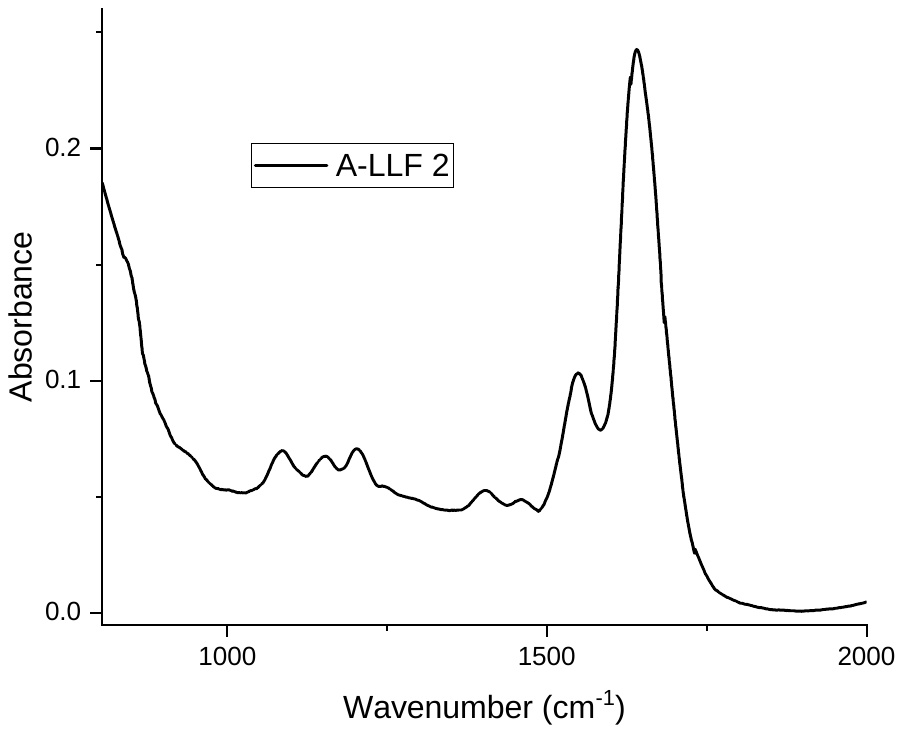

b)

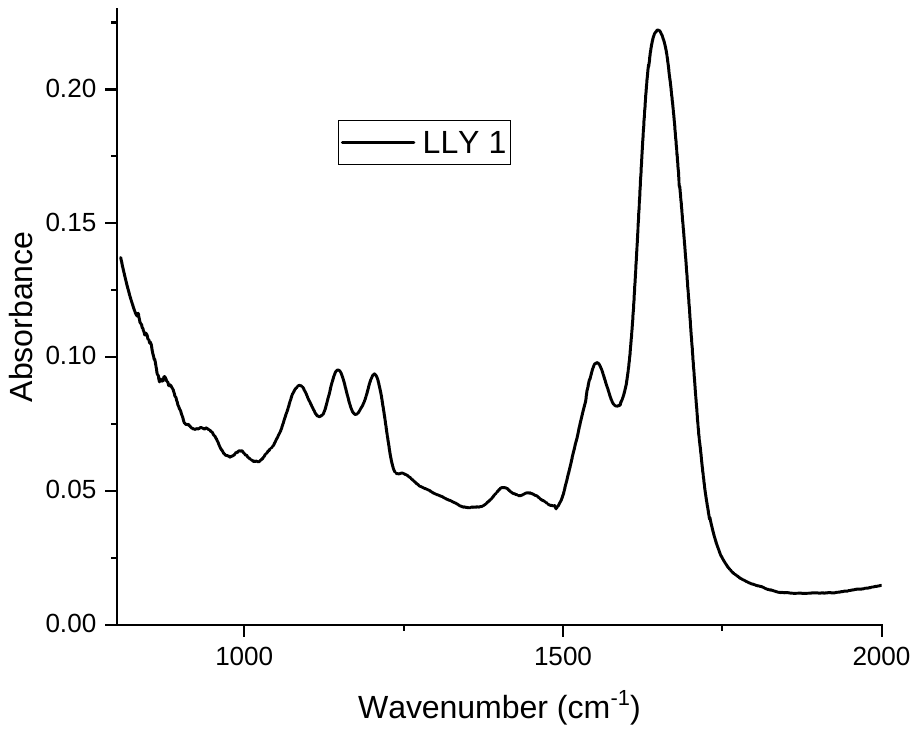

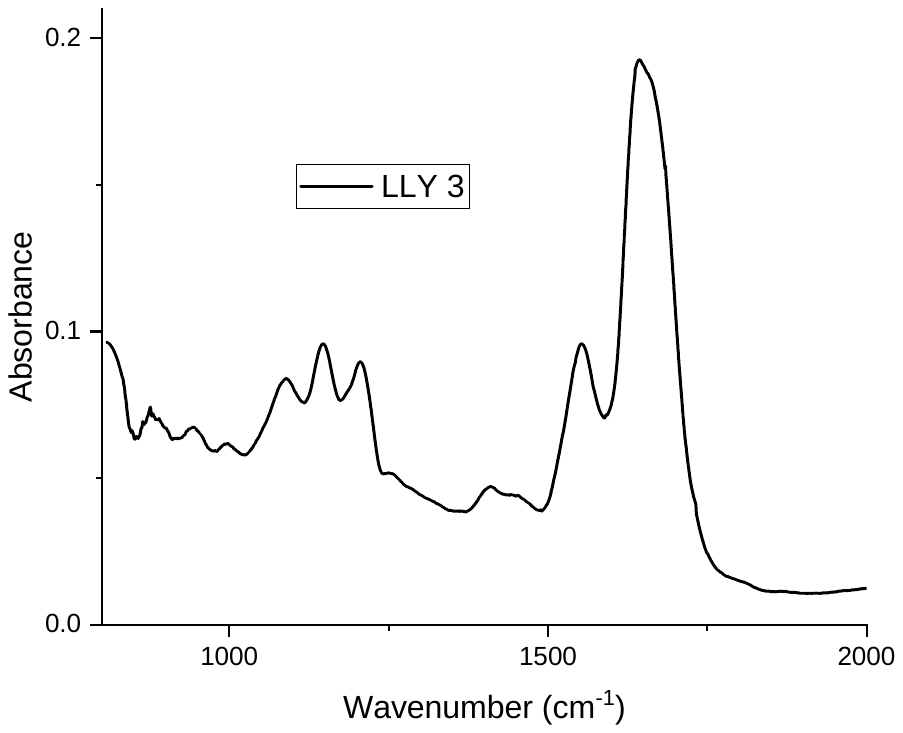

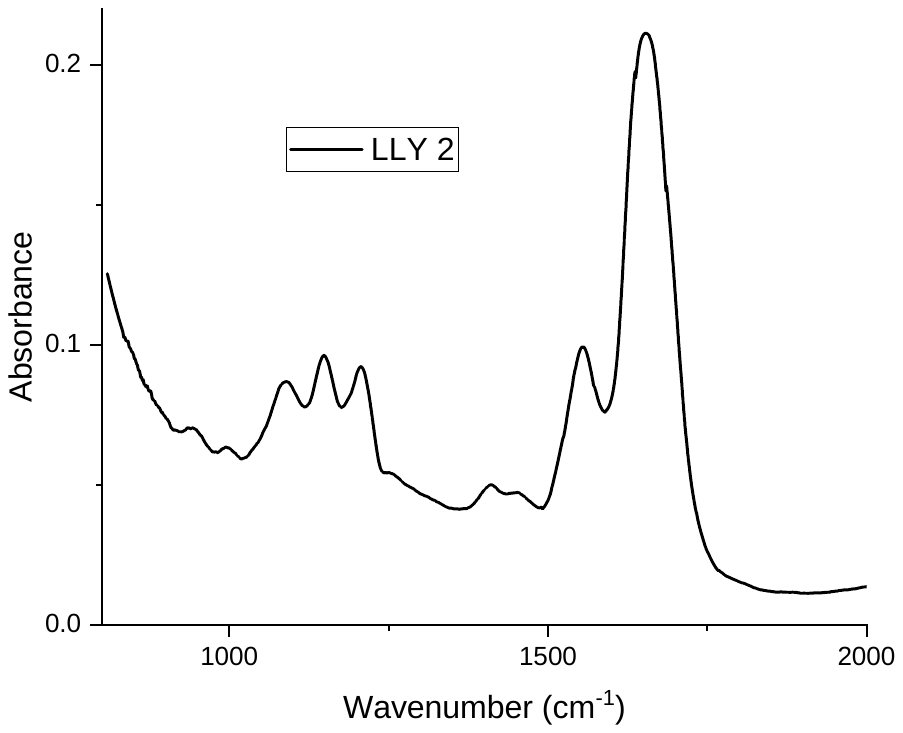

c)

d)

e)

**Figure S4.** Extended ATR-FTIR spectra measured in PBS containing 0.9 mM Ca^2+^ of 17-amino acid FP sequences of SARS-2 FP1+ (a); SARS-2 FP1+ Phe2Ala (b); FP1+ Phe8Tyr (c); FP1+ Leu6Ala (d); FP1+ Leu6Ser (e). The peptide backbone give rise to amide I (1600 – 1700 cm^-1^), corresponding to C=O stretching; amide II (1480 – 1575 cm^-1^), corresponding to N-H bending and C-N stretching; and amide III (1200 – 1350 cm^-1^), corresponding to C-N stretching, N-H bending, C-C stretching, and C-H bending. Of particular relevancy to peptide secondary structure are amide I and amide II bands. Since both C=O and N-H bonds are involved in hydrogen bonding, which gives insight into the secondary structure, amide I and amide II bands are generally used to yield information about peptide conformation (1,2). Among these amide bands, amide I is most widely used to determine peptide secondary structure as a result of its strong signal. Three individual spectra were collected for each FP and averaged to yield the spectra shown in Figure 5c of the main text.

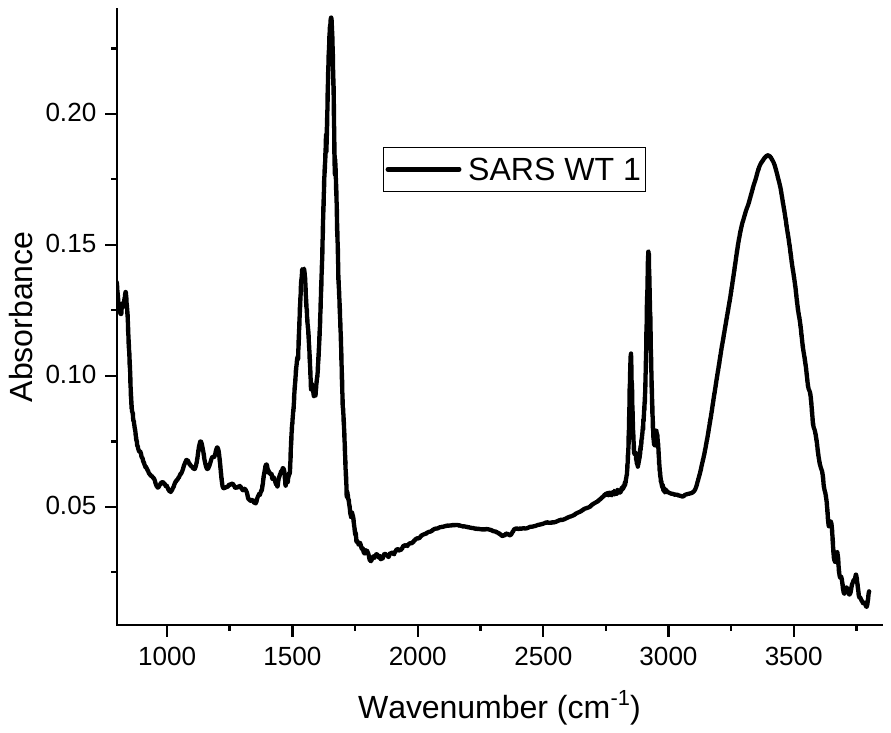

**Figure S5.** Representative ATR-FTIR spectrum encompassing peaks of C-H stretching, which represent the non-polar monolayer composed of 1-dodecanethiol (2800 – 3000 cm^-1^). The spectrum also includes peaks of O-H stretching of bulk PBS (3100 – 3750 cm^-1^).

**Relationship between peak positions of SARS-2 FP1 and MERS FP1 Amide I peaks (cm^-1^) and their corresponding adhesion forces (nN) in 60 vol % MeOH in PBS (corresponding to Tables 1 and 2 in main text).**

The results reveal a strong correlation between the conformation of the adsorbed FP1 peptides and the strength of their hydrophobic interaction. We note, however, that differences exist between the design of the AFM and ATR-FTIR experiments. In our AFM experiments, the FP sequences were covalently tethered via terminal thiol groups onto monolayers of tetraethylene glycol and then brought into contact with the non-polar surface of the AFM tip. In contrast, the ATR-FTIR experiments involve the adsorption of the FPs from bulk solution onto the non-polar surface of ATR-FTIR crystal. It is possible, therefore, that the conformational states of the peptides in the two experiments are not identical due to constraints on conformational degrees of freedom associated with covalent immobilization of the FP peptides used in our AFM experiments. Although we cannot rule out a role for these differences in our experiments, the strength of the correlation observed in Figure 5e between the Amide I peak wavenumber (i.e., peptide conformation) and the magnitude of hydrophobic pull-off force (Figure 5e) strongly suggests that these differences do not dominate our conclusions. We interpret our results to indicate that a single amino acid substitution at position 2, from Ala to Phe, profoundly influences the FP1 secondary structure, thereby changing the nanoscopic pattern in which the residues are presented during hydrophobic interaction with non-polar surfaces.

**Investigation of the secondary structure of the 17-amino acid FP1+ sequences with LLF substitutions in bulk PBS containing 0.9 mM Ca^2+^ and PBS containing 60 vol % MeOH.**

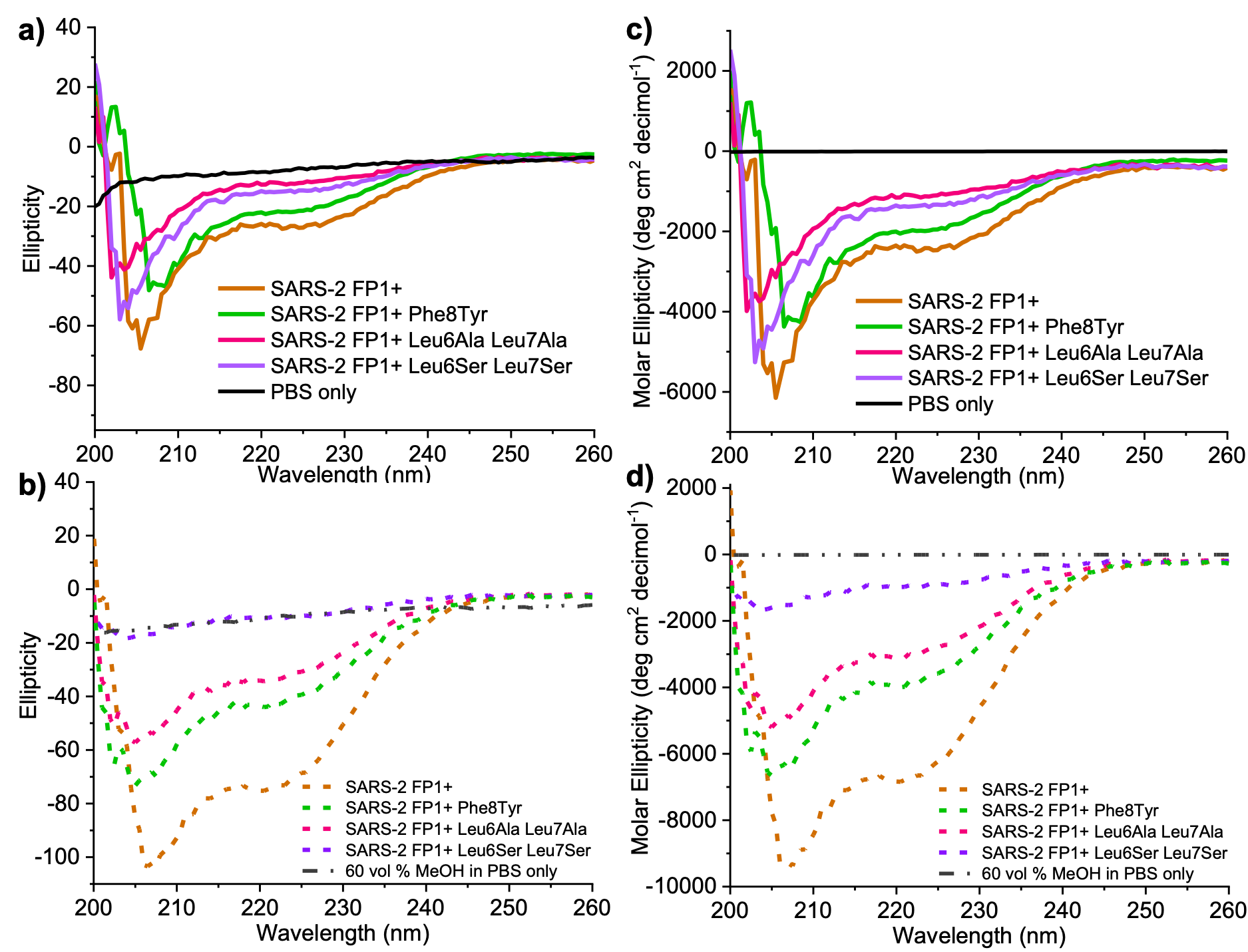

**Figure S6**. Circular dichroism of spectra the 17-amino acid FP1+ sequences with LLF substitutions in bulk PBS containing 0.9 mM Ca^2^ measured in ellipticity in units of mdeg (a) and converted to mean residue ellipticity in units of deg cm^2^ decimol^-1^ (c). CD spectra were measured in 60 vol % methanol in PBS containing 0.9 mM Ca^2^ in ellipticity (b) and mean residue ellipticity (d). All CD curves are the average of three independently collected spectra.

We used CD to investigate the secondary structure of SARS-2 FP1+ and its corresponding 17-amino acid sequences containing point substitutions to non-polar residues of LLF (Figure 6b). In bulk PBS at pH 7.4, the wild-type sequence exhibits a minimum at 208 nm and a weak signal around 222 nm, indicating a mix of random coil and α-helical conformational states (Figure 6b, brown spectrum). Substitution of LLF for polar (less non-polar) amino acids led to spectra suggestive of increasing random coil content. These results indicate that in bulk solution, substitution of LLF for less non-polar residues leads to conformational states that are distinct from the wild-type, suggesting the necessary cooperation between Phe 2 and LLF to induce greater α-helical content. Finally, greater α-helical content was found with the wild-type sequence in which Phe 2 flanks LLF, reinforcing the cooperative influence of Phe 2 and LLF in dictating the secondary structure of the FPs even before the presentation of the sequence to the non-polar surface.

Upon the addition of methanol, all FP sequences appear to adopt an increase in α-helical content (Figure 6b in main text; Figure S6b) compared to their respective secondary structure in PBS (Figure 6a in main text; Figure S6a). This suggests the strengthening of intramolecular hydrogen bonds within the peptides in their monomeric state, thus inducing α-helicity. Further, Phe 2-containing sequences exhibit more α-helical content than Ala 2-containing sequences, indicated by more negative spectroscopic peaks at 208 nm and 222 nm.

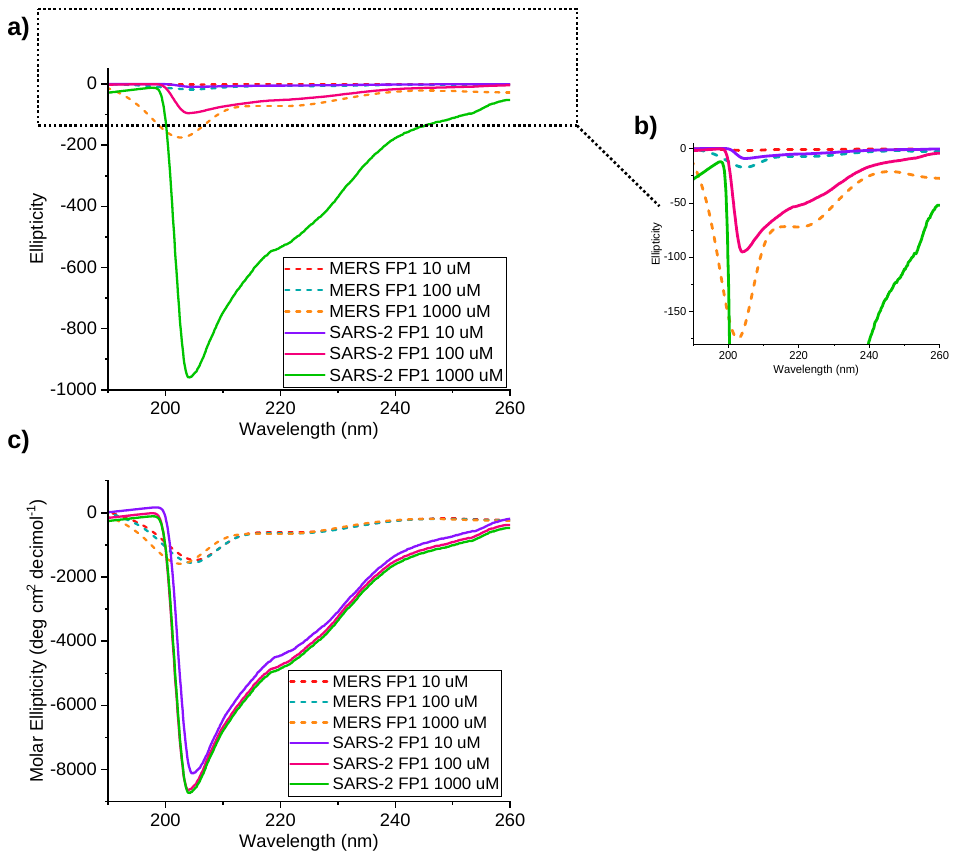

**Figure S7**. Circular dichroism of spectra SARS-2 FP1 (solid curves) and MERS FP1 (dashed curves) bulk PBS containing 0.9 mM Ca^2+^ in ellipticity in units of mdeg (a) and zoomed-in inset (b). The spectra were normalized to mean residue ellipticity in units of deg cm^2^ decimol^-1^ (c). All CD curves are the average of three independently collected spectra.

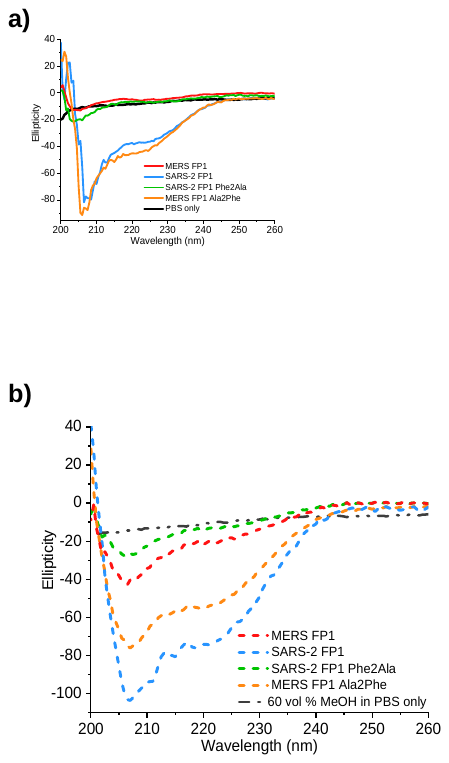

**Figure S8**. Circular dichroism of spectra the 11-amino acid FP1 sequences in ellipticity (units of mdeg) measured in bulk PBS containing 0.9 mM Ca^2+^ (a) and upon addition of 60 vol % MeOH to PBS containing 0.9 mM Ca^2+^ (b). These spectra correspond to Figure 6 in the main text. All CD curves are the average of three independently collected spectra.

|  | **Adhesive Force in PBS (nN)** | **Standard Deviation** | **Number of test events (n)** |
| --- | --- | --- | --- |
| **1.** MERS FP1 | 0.25 | 0.069 | 3240 |
| **2.** SARS-2 FP1 | 0.71 | 0.080 | 3109 |
| **3.** MERS FP1 Ala2Phe | 1.00 | 0.084 | 3822 |
| **4.** SARS-2 FP1 Phe2Ala | 0.23 | 0.015 | 3517 |
| **5.** SARS-2 FP1+ | 0.82 | 0.035 | 3221 |
| **6.** SARS-2 FP1+ Phe2Ala | 0.29 | 0.047 | 3481 |
| **7.** SARS-2 FP1+ Phe8Tyr | 0.60 | 0.028 | 3004 |
| **8.** SARS-2 FP1+ Leu6Ala | 0.50 | 0.033 | 3190 |
| **9.** SARS-2 FP1+ Leu6Ser | 0.30 | 0.018 | 3801 |

**Table S1.** Summary of adhesive forces in PBS, standard deviation, and number of test events (n) corresponding to the FP sequence used in performing *t*-tests. This table contains all FPs used in the study. The number of test events (n) correspond to the number of pull-off curves measured across 6 independent samples measured for each FP sequence in PBS.

Comparisons for t-tests:

I. 1 vs. 2 (SARS-2 FP1 vs. MERS FP1)
II. 1 vs. 4 (MERS FP1 vs. SARS-2 FP1 Phe2Ala)
III. 2 vs. 3 (SARS-2 FP1 vs. MERS FP1 Ala2Phe)
IV. 5 vs. 6 (SARS-2 FP1**+** vs. SARS-2 FP1**+** Phe2Ala)
V. 2 vs. 5 (SARS-2 FP1 vs. SARS-2 FP1**+**)
VI. 4 vs. 6 (SARS-2 FP1 Phe2Ala vs. SARS-2 FP1**+** Phe2Ala)
VII. 5 vs. 7 vs. 8 vs. 9 (SARS-2 FP1**+** vs. SARS-2 FP1+ Phe8Tyr vs. SARS-2 FP1+ Leu6Ala vs. SARS-2 FP1+ Leu6Ser)

| **Comparison** | **T value** | **Threshold value** | **p-value** |
| --- | --- | --- | --- |
| SARS-2 FP1 vs. MERS FP1 | 10.67 | 1.86 | < 0.05 |
| MERS FP1 vs. SARS-2 FP1 Phe2Ala | 0.69 | 1.86 | > 0.05 |
| SARS-2 FP1 vs. MERS FP1 Ala2Phe | 6.12 | 1.86 | < 0.05 |
| SARS-2 FP1+ vs. SARS-2 FP1+ Phe2Ala | 22.15 | 1.86 | < 0.05 |
| SARS-2 FP1 vs. SARS-2 FP1+ | 3.08 | 1.86 | < 0.05 |
| SARS-2 FP1 Phe2Ala vs. SARS-2 FP1+ Phe2Ala | 2.98 | 1.86 | < 0.05 |
| SARS-2 FP1+ vs. SARS-2 FP1+ | 12.02 | 1.86 | < 0.05 |

**Table S2.** Summary of *t*-test results for comparisons between various FPs discussed in the main text. With the exception of the comparison between MERS FP1 vs. SARS-2 FP1 Phe2Ala, we found that all p-values were < 0.05, revealing that the mean adhesive forces measured in PBS attributed to hydrophobic interactions were statistically different at a significance level of 95%. A p-value > 0.05 for MERS FP1 vs. SARS-2 FP1 Phe2Ala indicates that their mean force in PBS were statistically identical at a significance level of 95%.

| **Dulbecco’s Phosphate-Buffered Saline, 1X with calcium and magnesium** | |
| --- | --- |
| **Inorganic Salts** | **g/L** |
| CaCl_2_ (anhydrous) | 0.10 |
| KCl | 0.20 |
| KH_2_PO_4_ | 0.20 |
| MgCl_2_ • 6H_2_O | 0.10 |
| NaCl | 8.00 |
| Na_2_HPO_4_ • 7H_2_O | 2.1716 |

**Table S3.** Breakdown of components within PBS containing calcium and magnesium used in AFM, ATR-FTIR, and CD experiments in the main text.
